# LETR1 is a lymphatic endothelial-specific lncRNA that governs cell proliferation and migration through KLF4 and SEMA3C

**DOI:** 10.1101/2020.05.25.114546

**Authors:** Luca Ducoli, Saumya Agrawal, Eliane Sibler, Tsukasa Kouno, Carlotta Tacconi, Chung-Chao Hon, Simone D. Berger, Daniela Müllhaupt, Yuliang He, Marco D’Addio, Lothar C. Dieterich, Piero Carninci, Michiel J. L. de Hoon, Jay W. Shin, Michael Detmar

**Author notes:** These two authors share co-senior authorship (Jay W. Shin, Michael Detmar). Correspondence: Prof. Dr. Michael Detmar, Institute of Pharmaceutical Sciences, ETH Zurich, Vladimir-Prelog-Weg 3, 8093 Zurich, Switzerland.

## Abstract

Recent studies have revealed the importance of long noncoding RNAs (lncRNAs) as tissue-specific regulators of gene expression. There is ample evidence that distinct types of vasculature undergo tight transcriptional control to preserve their structure, identity, and functions. We determined, for the first time, the global lineage-specific lncRNAome of human dermal blood and lymphatic endothelial cells (BECs and LECs), combining RNA-Seq and CAGE-Seq. A subsequent genome-wide antisense oligonucleotide-knockdown screen of a robust set of BEC- and LEC-specific lncRNAs identified LETR1 as a critical gatekeeper of the global LEC transcriptome. Deep RNA-DNA, RNA-protein, and phenotype rescue analyses revealed that LETR1 acts as a nuclear *trans*-acting lncRNA modulating, via key epigenetic factors, the expression of essential target genes, including KLF4 and SEMA3C, governing the growth and migratory ability of LECs. Together, our study provides new evidence supporting the intriguing concept that every cell type expresses precise lncRNA signatures to control lineage-specific regulatory programs.

## Introduction

The blood and lymphatic vascular systems are essential for the efficient transport of oxygen, nutrients, signaling molecules, and leukocytes to and from peripheral tissues, the removal of waste products, and the preservation of fluid homeostasis. Increased activation or impaired function of these vascular networks represent a hallmark of many pathological conditions, including cancer progression, chronic inflammatory diseases, and diseases leading to blindness^1-3^.

During development, the blood vascular system arises from endothelial cell progenitors that differentiate from mesodermal cells, mostly through the expression of the transcription factor (TF) ETV2. Activation of the VEGFA/VEGFR2 signaling and expression of blood vascular endothelial cell (BEC) markers, such as NRP1 and EphrinB2, further differentiate these precursor cells into BECs, which then form the hierarchical network of blood vessels^4^. In contrast, lymphatic vasculogenesis starts after the establishment of the blood circulatory system. Thereafter, a distinct subpopulation of endothelial cells lining the cardinal vein starts differentiating by expressing the TF PROX1, the master regulator of lymphatic endothelial cell (LEC) identity, via the TFs SOX18 and COUPTFII. Once exiting the veins, LECs starts expressing other lymphatic-specific markers, such as podoplanin, VEGFR3, and NRP2, and they migrate, in a VEGFC-dependent manner, to form the primary lymph sacs from which the lymphatic vascular system further develops following sprouting, branching, proliferation, and remodeling processes^5^. However, a nonvenous origin of LECs has also been described in the skin, mesenteries, and heart^6-8^. In adulthood, while the blood and lymphatic vasculature are generally quiescent, they can be readily activated in pathological conditions such as wound healing, inflammation, and cancer by disturbance of the natural balance of pro- and anti-(lymph)angiogenic factors^1,9^. Therefore, this complex regulatory network requires precise control of gene expression patterns at both transcriptional and post-transcriptional levels in order to ensure proper maturation, differentiation, and formation of blood and lymphatic vessels.

In this scenario, many studies have recently revealed the importance of a new member of the noncoding RNA clade, termed long noncoding RNAs (lncRNAs), in the regulation of gene activity^10,11^. In particular, the FANTOM (functional annotation of the mammalian genome) consortium pioneered the discovery of the noncoding RNA world by providing, through Cap analysis of gene expression (CAGE-Seq), the first evidence that large portions of our genome are transcribed, producing a multitude of sense and antisense transcripts^12^. In the latest genome annotation, lncRNAs, which are arbitrarily defined as noncoding RNAs longer than 200 nucleotides, constitute approximately 72% of the transcribed genome^13^, whereas mRNAs comprise only 19%, indicating the need for functional annotation of lncRNAs. Importantly, lncRNAs have recently been shown to display a higher tissue-specificity than mRNAs, suggesting them as new players in the regulation of cell-type-specific gene expression programs^14^.

Since lncRNAs lack a protein-coding role, their primary categorization is based on their genomic location and orientation relative to protein-coding genes^15^. lncRNAs can reside either between protein-coding genes (intergenic, lincRNAs), between two exons of the same gene (intronic lncRNAs), antisense to protein-coding transcripts (antisense lncRNAs), or in promoters and enhancers (natural antisense transcripts or transcribed from bidirectional promoters)^16-18^. lncRNAs may regulate gene expression through a multitude of mechanisms depending on their subcellular localization. For instance, in the nucleus, lncRNAs can act as a scaffold for TFs, chromatin remodeling complexes, or ribonucleoprotein complexes (RNPs), indicating a potential role in transcriptional regulation^19^. Nuclear lncRNAs can furthermore act in *cis* or *trans* to regulate gene expression by the recruitment of activating and repressive epigenetic modification complexes. *Cis*-acting lncRNAs, such as the 17-kb X chromosome-specific transcript Xist, regulate gene expression of adjacent genes by directly targeting and tethering protein complexes^20,21^. On the other hand, *trans*-acting lncRNAs, such as the HOTAIR lncRNA, regulate gene expression at distinct genomic loci across the genome by serving as a scaffold that assists the assembly of unique functional complexes^22^.

In blood vessels, some lncRNAs have been reported to play a role in angiogenesis (MALAT-1, lnc-Ang362)^23-25^, tumor-induced angiogenesis (MVIH, HOTAIR)^26,27^, and proliferation as well as cell junction regulation of endothelial cells (MALAT-1, Tie-1AS)^24,28^. In contrast, while cancer cell expression of the antisense noncoding RNA in the INK4 locus (ANRIL) and of the lymph node metastasis associated transcript 1 (LNMAT1) have been associated with lymphangiogenesis and lymphatic metastasis^29,30^, lymphatic endothelial-specific lncRNAs have not been identified or functionally characterized so far.

In the context of the international FANTOM6 project, which aims to functionally annotate all lncRNAs present in our genome, we first determined lineage-specific lncRNAs associated with human primary dermal LECs and BECs by combining RNA-Seq and CAGE-Seq analyses. Genome-wide functional interrogation after antisense-oligonucleotide (ASO) knockdown of robustly selected LEC and BEC lncRNAs, allowed us to identify LINC01197, which we renamed LETR1 (lymphatic endothelial transcriptional regulator), as a lymphatic endothelial-specific lncRNA that functions in the transcriptional regulation of LEC growth and migration. We demonstrated that LETR1 is a *trans*-acting lncRNA that acts as a protein scaffold in order to facilitate the assembly of unique functional epigenetic complexes involved in gene expression regulation. Through these interactions, LETR1 controls intricated transcriptional networks to fine-tune the expression, above all, of essential proliferation- and migration-related genes, including the tumor-suppressor TF KLF4 and the semaphorin guidance molecule SEMA3C.

## Results

### Identification of a core subset of vascular lineage-specific lncRNAs

To identify vascular lineage-specific lncRNAs, we performed both RNA-Seq and CAGE-Seq^31^ of total RNA isolated from neonatal human primary dermal LECs and BECs (Supplementary Figure 1a, b). Compared to RNA-Seq, CAGE-Seq allows mapping transcription start sites (TSSs) after quantification of the expression of 5’-capped RNAs^32^. To ensure endothelial cell specificity, we included RNA-Seq and CAGE-Seq data from neonatal human primary dermal fibroblasts (DFs)^33^. In a first step, we performed differential expression (DE) analysis of RNA-Seq and CAGE-Seq of LECs against BECs, LECs against DFs, and BECs against DFs using EdgeR^34^. From defined LEC- or BEC-specific genes (see Methods section), we selected genes annotated as lncRNAs in the recently published FANTOM CAT database^13^. Finally, we overlapped the RNA-Seq and CAGE-Seq results to select lncRNAs identified as differentially expressed using both techniques (Figure 1a). RNA-Seq identified 832 LEC- and 845 BEC-associated lncRNAs, after the exclusion of 232 LEC and 672 BEC lncRNAs also expressed in DFs (Figure 1b). In contrast, CAGE-Seq identified 277 LEC lncRNAs and 243 BEC lncRNAs, after the removal of 143 BEC and 282 LEC lncRNAs also expressed in DFs (Figure 1c). The intersection of these two methods revealed 142 LEC and 160 BEC lncRNAs specifically expressed in either LECs or BECs. We defined these subsets as LEC and BEC core lncRNAs (Figure 1d and Supplementary Table 1).

**Figure 1:**
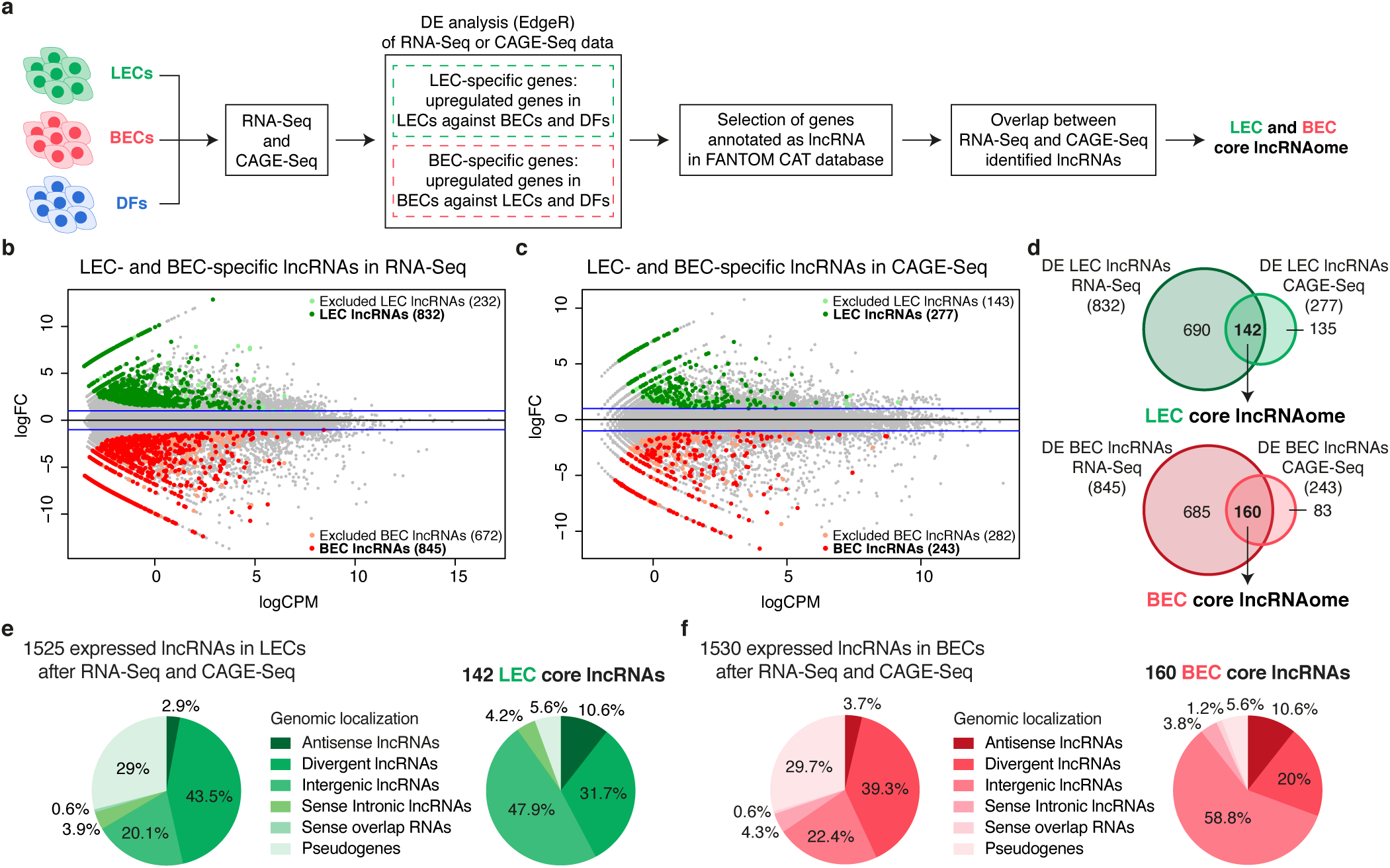
Identification of a core subset of vascular lineage-specific lncRNAs. **(a)** Schematic representation of the analysis pipeline. Total RNA was extracted from two replicates of neonatal LECs and BECs derived from the same donor and subjected to both RNA-Seq and CAGE-Seq. Neonatal dermal Fibroblasts (DFs) data from FANTOM6 database^33^ were included to increase endothelial cell specificity. After differential expression (DE) analysis of RNA-Seq and CAGE-Seq using EdgeR^34^, we overlapped the results from RNA-Seq and CAGE-Seq to select lncRNAs differentially expressed in both techniques. **(b, c)** MA plots displaying log2FC against expression levels (logCPM) of DE genes after RNA-Seq (b) and CAGE-Seq (c) between LECs and BECs. Green dots represent LEC-specific lncRNAs, and red dots represent BEC-specific lncRNAs (FDR < 0.01). Blue horizontal lines show the chosen |log2FC| > 1. Light green and light red dots represent the lncRNAs excluded from the analysis because also expressed in DFs. **(d)** Venn diagrams showing the overlap between RNA-Seq and CAGE-Seq and the identified LEC- (top) and BEC (bottom) core lncRNAs. LEC and BEC core lncRNAs are listed in Supplementary Table 1. **(e, f)** Pie charts showing the genomic classification according to FANTOM CAT database^13^ of LEC (e) and BEC (f) core lncRNAs compared to lncRNAs generally expressed in LECs or BECs by both RNA-Seq and CAGE-Seq (RNA-Seq: TPM > 0.5 and CAGE-Seq: CPM > 0.5).

To characterize the identified LEC and BEC core lncRNA subsets, we analyzed their genomic classification related to protein-coding genes, using the FANTOM CAT^13^ annotations. We found that the largest fraction of both LEC and BEC core lncRNAs were categorized as intergenic lncRNAs (47.9% for LEC and 58.8% for BEC), with a significant enrichment compared to all expressed lncRNAs (fold enrichment = 1.9 resp. 2.17, P-value < 0.05) (Figure 1e, f). Gene ontology (GO) analysis of lncRNAs flanking protein-coding genes using Genomic Regions Enrichment of Annotations Tool (GREAT)^35^ and g:Profiler^36^ showed that both core lncRNA subsets mainly reside near genes related to vascular development, tissue morphogenesis, and endothelial cell function, including proliferation, migration, and adhesion (Supplementary Figure 1c-f). These results are intriguing since several intergenic lncRNAs have previously been reported to play a prominent role in the regulation of gene expression in a cell-specific manner ^11^.

### Identification of lncRNA candidates for functional characterization by antisense oligonucleotides (ASOs)

To further select lncRNA candidates for genome-wide functional screening, we relied on the FANTOM CAT annotations^13^. Firstly, we filtered for lncRNAs with a conserved transcription initiation region (TIR) and/or exon regions, based on overlap with predefined genomic evolutionary rate profiling (GERP) elements^37^. Secondly, we selected for actively-transcribed lncRNAs with an overlap between TSSs and DNase hypersensitive sites (DHSs). Thirdly, filtering for expression levels in LEC and BEC RNA-Seq and CAGE-Seq datasets (see Methods section) led to the identification of 5 LEC and 12 BEC lncRNAs that are potentially conserved at the sequence level, actively transcribed, and robustly expressed in the respective endothelial cell types (Figure 2a-c). Finally, we identified through qPCR 2 LEC (AL583785.1 and LETR1), and 2 BEC (LINC00973 and LINC01013) lncRNAs that were consistently differentially expressed between LECs and BECs derived from newborn and adult skin samples (Figure 2d, e).

**Figure 2:**
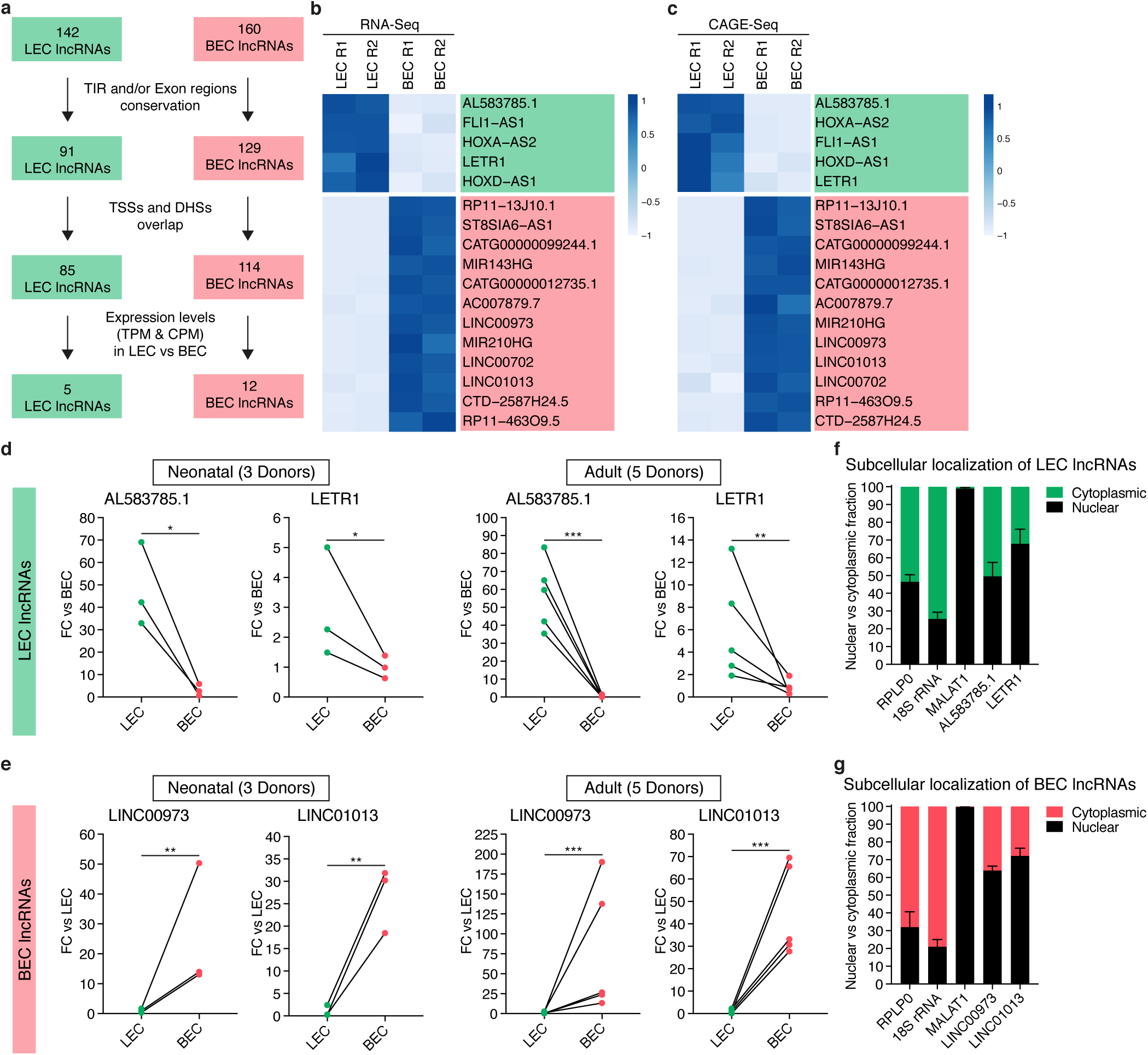
Identification of lncRNA candidates for functional characterization by antisense oligonucleotides (ASOs). **(a)** Diagram showing the criteria applied to select the final LEC and BEC lncRNA candidates. First criterion: sequence conservation of transcription initiation regions (TIR) and/or exon regions. Second criterion: overlap between transcription start sites (TSSs) and DNase hypersensitive sites (DHSs) as a hint for active transcription. Third criterion: expression level cut-offs between LECs and BECs. For LEC lncRNAs: TPM and CPM in BECs < 5; TPM and/or CPM > 10 in LECs. For BEC lncRNAs: TPM and CPM in LECs < 5; TPM and CPM > 10 in BECs. **(b, c)** Heat maps based on expression levels of RNA-Seq (b, TPM, 2 replicates) and CAGE-Seq (c, CPM, 2 replicates) of 5 LEC (light green) and 12 BEC (light red) lncRNAs filtered from a. Color code for row Z-Score values on a scale from −1 to +1. Genes were ordered by log2FC values of RNA-Seq or CAGE-Seq (c). **(d, e)** Validation of the differential expression of 2 LEC (d) and 2 BEC (e) target lncRNAs in LECs and BECs derived from 3 neonatal and 5 adult donors. qPCR results are displayed as fold change (FC) against average BEC or LEC expression as mean + SD. GAPDH was used as the housekeeping gene. *P < 0.05, **P < 0.01, ***P < 0.001 using paired two-tailed Student’s t-test on ΔCt values against BEC or LEC. **(f, g)** Determination of the LEC (f) and BEC (g) lncRNAs subcellular localization through cellular fractionation followed by qPCR in neonatal LECs and BECs derived from 3 donors. Bars represent percentages nuclear (black) and cytoplasmic (green for LEC, red for BEC) fraction displayed as mean + SD.

Since antisense oligonucleotide (ASO) GapmeRs are more effective in reducing the expression of nuclear lncRNAs than short interference RNAs (siRNAs)^38,39^, we analyzed the subcellular localization of the 2 LEC and 2 BEC lncRNAs in LECs and BECs, using cellular fractionation followed by qPCR. For LEC lncRNAs, AL583785.1 was almost equally distributed between cytoplasm and nucleus, whereas LETR1 showed a higher nuclear distribution (Figure 2f). Both LINC00973 and LINC01013 were mainly localized in the nucleus (Figure 2g). Therefore, we next used the ASO-based approach to analyze the genome-wide transcriptional changes upon knockdown of the 2 LEC and 2 BEC lncRNAs. After testing their knockdown efficiencies, we selected three out of five ASOs for each lncRNA target (Supplementary Figure 2a-d and Supplementary Table 2).

### Transcriptional profiling after LETR1-ASOKD indicates potential functions in cell growth, cell cycle progression, and migration of LECs

To investigate the potential functional relevance of the 2 LEC and 2 BEC lncRNAs, we first transfected LECs and BECs with three independent ASOs per target, followed by CAGE-Seq (Figure 3a and Supplementary Figure 3a, b). Next, we performed DE analysis by comparing the combined results of the ASOs per target with their scramble controls, using EdgeR with a generalized linear model (GLM)^34^. Finally, we defined DE genes by a false discovery rate (FDR) < 0.05 and a log2 fold change (log2FC) > 0.5 resp. < −0.5 (Supplementary Table 3). We found that ASO knockdown (ASOKD) of AL583785.1 in LECs and of LINC00973 and LINC01013 in BECs showed rather modest changes in gene expression. AL583785.1-ASOKD caused changes of only 9 genes (4 up and 5 down), LINC00973-ASOKD of 43 genes (6 up and 37 down), and LINC01013-ASOKD of 24 genes (2 up and 22 down) (Supplementary Figure 3c-g).

**Figure 3:**
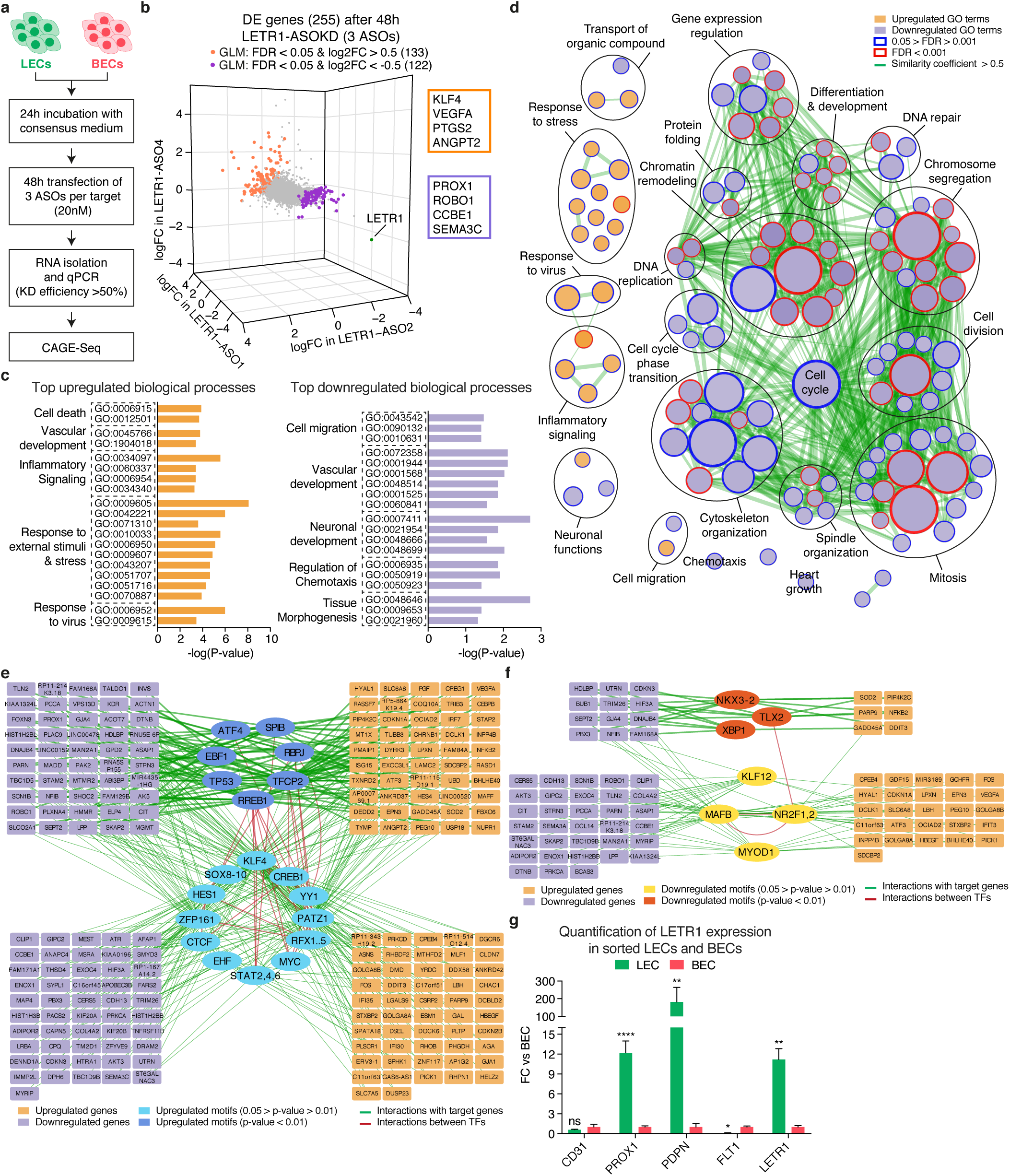
Transcriptional profiling after LETR1-ASOKD indicates potential functions in cell growth, cell cycle progression, and migration of LECs. **(a)** Schematic representation of the ASO-mediated perturbation strategy of two replicates of neonatal LECs and BECs derived from the same donor. Only samples with ASOKD efficiency > 50% were subjected to CAGE-Seq. **(b)** 3-dimensional scatter plot showing log2FC values calculated between single ASO against LETR1 and control ASO using EdgeR^34^. Orange and purple dots represent significantly (FDR < 0.05) up- and downregulated genes (|logFC| > 0.5) after applying a generalized linear model (GLM) design. Green dot shows LETR1. Orange and purple text boxes show examples of DE genes with important functions in vascular biology. **(c)** Top significantly (P-value < 0.05) enriched GO terms for biological processes of selected up- and downregulated genes after LETR1-ASOKD, using g:ProfileR^36^ (relative depth 2-5). Terms were manually ordered according to their related biological meaning. Enriched GO terms after g:ProfileR are listed in Supplementary Table 4. **(d)** Network of significantly (FDR < 0.05) enriched biological processes in LETR1-ASOKD data after GSEA analysis^45^ generated using Cytoscape and Enrichment Map^93^. Orange and purple nodes represent up- and downregulated GO terms, respectively. Node size represents the total number of genes included in the gene sets. Color code of node borders represents FDR values on a scale from 0.05 (white) to 0.001 (red). Edge width represents the portion of shared genes between gene sets starting from 50% similarity. To improve visualization, related biological processes were grouped manually using Wordcloud. Enriched GO terms after GSEA analysis are listed in Supplementary Table 5. **(e, f)** MARA network of up- (e) and downregulated (f) transcription factor binding motifs and their connection with 255 DE genes after LETR1-ASOKD. Only genes with at least a connection are displayed. Cyan/dark yellow ellipses are differentially motifs with P-value between 0.05 and 0.01, and blue/brown ellipses are differentially motifs with P-value < 0.01. Orange and purple rectangles correspond to up- and downregulated genes, respectively. Green edges reflect the connections between deregulated genes and active motifs. Red edges represent the connections between motifs. Enriched TF motifs are listed in Supplementary Table 6. **(g)** Expression levels of LETR1 and endothelial (CD31), lymphatic (PROX1, PDPN), and blood (FLT1) markers in freshly sorted LECs and BECs derived from 3 donors of healthy human skin samples. Bars represent FC values against BEC displayed as mean ± SD. GAPDH was used as the housekeeping gene. *P < 0.05, **P < 0.01, ****P < 0.0001, ns not significant using paired two-tailed Student’s t-test on ΔCt values against BEC.

In contrast, ASOKD of LETR1 had a high impact on the global transcriptome of LECs, resulting in 133 up- and 122 downregulated genes (Figure 3b and Supplementary Figure 3f). Among these, several genes have previously been reported to play prominent roles in vascular development and differentiation pathways, including PTGS2, KLF4, VEGFA, and ANGPT2 among the upregulated genes, and PROX1, CCBE1, SEMA3C, and ROBO1 among the downregulated genes^40-44^. GO analysis for biological processes using g:ProfileR^36^ revealed that indeed both up- and downregulated genes were enriched (P-value < 0.05) for terms related to vascular development. In addition, upregulated genes were mainly involved in cell death, inflammatory signaling, and response to external stimuli, whereas downregulated genes were primarily related to the regulation of cell migration and chemotaxis (Figure 3c and Supplementary Table 4). Gene set enrichment analysis (GSEA)^45^ also identified significant (FDR < 0.05) biological processes related to cell migration, chemotaxis, and response to external stimuli/virus. More importantly, several downregulated biological processes were associated with cell growth, cell cycle progression, and cytoskeleton organization (Figure 3d and Supplementary Table 5).

To identify TFs potentially affected by LETR1-ASOKD, we performed motif activity response analysis (MARA)^46^ by analyzing the activity of 348 regulatory motifs in TF sites in the proximal promoters of highly expressed genes in knockdown and control samples (see Methods section). We found 19 upregulated and 7 downregulated motifs, among which were binding sites related to several TFs known to be essential for LEC biology, including STAT6, KLF4, NR2F2 (COUPTF-II), and MAFB^43,47,48^ (P-value < 0.05, Supplementary Table 6). Interestingly, KLF4 was the only TF to be also upregulated on the transcriptional level upon LETR1-ASOKD. Based on the MARA analysis, we next reconstructed a gene regulatory network with the 255 genes affected by LETR1-ASOKD. We identified modules of up and downregulated genes linked with the identified TF motifs. Among these modules, we found genes associated with endothelial cell proliferation and migration, such as VEGFA, MAFF, ANGPT2, RASD1, PROX1, SEMA3C, and ROBO1^40,49,50^ (Figure 3e, f). Overall, these results suggest that the absence of LETR1 had a critical impact on the global transcriptome of LECs by affecting complex TF regulatory networks targeting essential genes largely involved in endothelial cell differentiation, proliferation, and migration.

Before characterizing the biological and molecular function of LETR1 in LECs, we sought to validate its lymphatic endothelial cell-selective expression *in situ*. Hence, we analyzed the expression levels of LETR1 and specific blood and lymphatic markers in freshly isolated LECs and BECs from human skin samples, using flow cytometry followed by qPCR (Supplementary Figure 3h). As shown in Figure 2d, LETR1 was more highly expressed in LECs isolated *ex vivo* than BECs. Remarkably, we observed that the LEC specificity of LETR1 was more pronounced in freshly isolated ECs than cultured ECs, similar to the LEC lineage-specific TF PROX1 (Figure 3g).

### Knockdown of LETR1 reduces cell growth, cell cycle progression, and migration of LECs *in vitro*

To investigate the potential effects of LETR1-ASOKD on LEC growth, we performed cell growth assays based on dynamic imaging analysis. We found that LETR1-ASOKD strongly reduced cell growth of LECs over time (Figure 4a and Supplementary Figure 4a, b). To study whether the cell growth phenotype was not due to off-target effects of the ASOs, we also performed cell growth assays after CRISPR interference (CRISPRi)^51^. Consistently, we found that CRISPRi-KD of LETR1 also significantly reduced the growth rate of LECs. However, due to the lower knockdown efficiency, the effect was less prominent compared to ASOKD (Figure 4b and Supplementary Figure 4c, d). Next, we analyzed the cell cycle progression of LECs upon LETR1-ASOKD, using flow cytometry. Double staining for Ki-67 (proliferation marker) and propidium iodide (PI, DNA content) showed that LETR1-ASOKD significantly increased the percentage of LECs arrested in G0 (Figure 4c, d and Supplementary Figure 4e). Although there was a slight increase of subG0 LECs in LETR1-ASOKD samples, analysis of cleaved caspase 3-positive cells showed that LETR1-ASOKD did not consistently induce apoptosis in LECs, suggesting an alternative cell death pathway (Supplementary Figure 4f).

**Figure 4:**
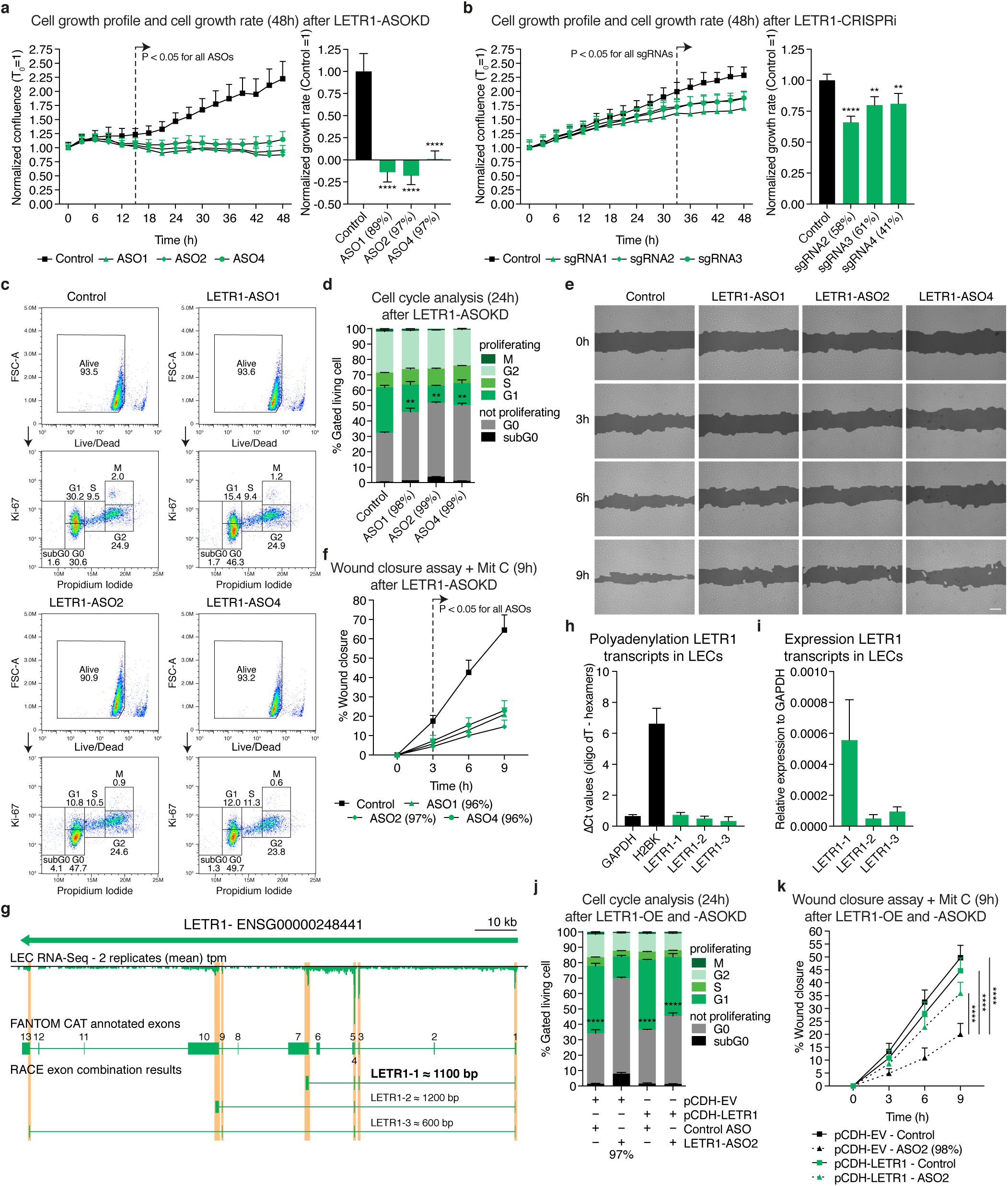
Knockdown of LETR1 reduces cell growth, cell cycle progression, and migration of LECs *in vitro*. **(a, b)** Cell growth profiles and cell growth rates of neonatal LECs over 48h after ASOKD (a) or CRISPRi-KD (b) of LETR1 using IncuCyte. Sample’s confluences were normalized to T_0_. Growth rates were calculated as the slope of linear regression and normalized to control ASO/sgRNA. **(c)** Representative flow cytometry plots of neonatal LECs after 24h LETR1-ASOKD. Cells were firstly gated with live/dead Zombie staining (upper plots). Resulting living cells were further gated for non-proliferating stages subG0 and G0, and proliferating stages G1, S, G2, and M, using propidium iodide (IP) and Ki-67 (lower plots). **(d)** Quantification of the cell cycle progression analysis of neonatal LECs after 24h LETR1-ASOKD. Bars represent percentages of gated living cells in subG0, G0, G1, S, G2, and M. Statistical analysis was performed on G0 populations. **(e)** Representative images of the wound closure assay (9h) in neonatal LECs after LETR1-ASOKD. Confluence mask is shown for all time points. Before scratch, cells were incubated for 2h with 2*µ*g/mL Mitomycin C (proliferation inhibitor) at 37°C. Scale bar represents 200*µ*m. **(f)** Quantification of the wound closure assay (up to 9h) of neonatal LECs after LETR1-ASOKD. Percentages were determined for each time point using TScratch^98^. **(g)** Schematic representation of 3’ RACE results depicting the three LETR1 transcripts expressed in LECs: LETR1-1 (approx. 1,100bp), LETR1-2 (approx. 1,200bp), LETR1-3 (approx. 600bp). RNA-Seq signal was visualized through the Zenbu genome browser^106^. LETR1 transcript sequences are listed in Supplementary Table 7. **(h)** Comparison of qPCR levels of GAPDH (polyA+), H2BK (polyA-), LETR1-1, LETR1-2, LETR1-3 after cDNA synthesis with either oligodT or random hexamers primers in neonatal LECs derived from 3 donors. **(i)** Expression of LETR1-1, LETR1-2, and LETR1-3 relative to housekeeping gene GAPDH in neonatal LECs derived from 3 donors. **(j)** Quantification of the cell cycle progression analysis of pCDH-empty vector (pCDH-EV) and pCDH-LETR1 infected neonatal LECs after 24h LETR1-ASOKD. Statistical analysis was performed on G0 populations. **(k)** Quantification of the wound closure assay (up to 9h) of pCDH-EV and pCDH-LETR1 infected neonatal LECs after LETR1-ASOKD. Data are displayed as mean + SD (n = 10 in a, f, and k; n = 5 in b; n = 3 in h, i, and j; n = 2 in d). Percentages represent LETR1 knockdown efficiencies after the experiments. **P < 0.01, ****P < 0.0001 using ordinary one-way (for a, b, d, and j) and two-way (for a, b, f, and k) ANOVA with Dunnett’s multiple comparisons test against control ASO/sgRNA or LETR1-ASO2 – control siRNA. All displayed *in vitro* assays were performed in neonatal LECs derived from the same donor.

Since the transcriptional studies also indicated a potential role of LETR1 in cell migration, we performed wound-closure assays (“scratch assays”) after LETR1-ASOKD in LECs. We observed a significant reduction of LEC migration compared to control (Figure 4e, f and Supplementary Figure 4g). Similarly, LETR1-ASOKD significantly inhibited LEC migration in a trans-well hapto-chemotactic assay (Supplementary Figure 4h, i).

We next studied whether ectopic overexpression of LETR1 could rescue the proliferation and migration phenotypes observed after knockdown of endogenous LETR1. We first determined the most abundant transcript variant of LETR1 in LECs, using 3’ rapid amplification of cDNA ends (3’ RACE) followed by qPCR. From the determined LETR1 TSS, we identified three primary polyadenylated LETR1 transcripts that overlapped with the RNA-Seq signal in LECs, where the LETR1-1 variant had the highest expression level based on 3’ RACE, RNA-Seq, and qPCR (Figure 4g-i and Supplementary Table 7). Subsequently, we overexpressed the most abundant transcript variant in LECs using a lentiviral vector and analyzed the cell cycle progression and cell migration after LETR1-ASOKD. We performed both assays with the most effective LETR1-ASO2, which binds to the first intron recognizing exclusively the endogenous but not ectopically overexpressed LETR1 (Supplementary Figure 2b). The reintroduction of ectopic LETR1 significantly ameliorated both phenotypes, further supporting the role of LETR1 in cell growth and migration regulation. Overexpression of LETR1 *per se* did not enhance both cellular functions compared to control, implicating a possible saturation of the regulatory system (Figure 4j, k and Supplementary Figure 4j-l).

### LETR1 is a nuclear lncRNA interacting *in trans* with DNA regions near a subset of differentially expressed genes

A first step to study the molecular mechanism of a lncRNA of interest is to analyze its subcellular distribution at the single-molecule level^52^. To this end, we performed single-molecule RNA fluorescence in situ hybridization (smRNA-FISH) in cultured LECs and human skin samples^53^. Consistent with the cellular fractionation data (Figure 2f), LETR1 was predominantly localized in the nucleus of cultured LECs, showing a broad nuclear distribution with distinct foci (Figure 5a, b). This localization pattern was also observed in lymphatic vessels in human skin (Supplementary Figure 5a), suggesting that LETR1 might exert a chromatin-related function *in vitro* as well as *in vivo*.

**Figure 5:**
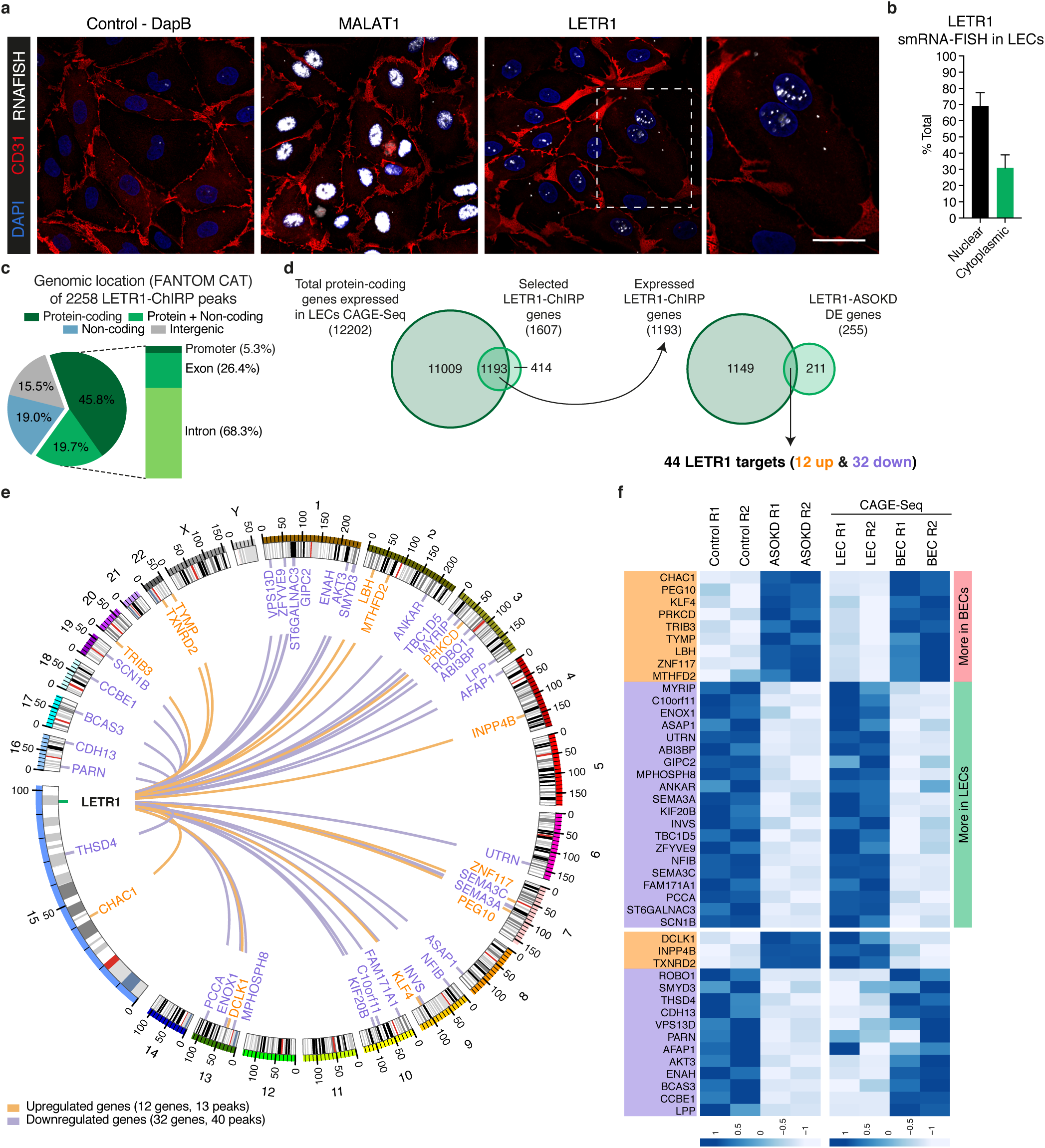
LETR1 is a nuclear lncRNA interacting *in trans* with DNA regions near a subset of differentially expressed genes. **(a)** Representative images of negative control dapB (bacterial gene), MALAT-1 (nuclear lncRNA), and LETR1 expression using smRNA-FISH. Immunostaining of endothelial cell marker CD31 was used to outline cell shape. Scale bars represent 20*µ*m. **(b)** Quantification of the nuclear (green) and cytoplasmic (black) smRNA-FISH signal of LETR1 in neonatal LECs derived from 2 donors quantified with ImageJ^97^. Bars represent nuclear and cytoplasmic percentages displayed as mean + SD. **(c)** Pie chart showing the genomic localization of 2,258 LETR1 peaks in protein-coding, overlap between protein-coding and noncoding, noncoding, and intergenic regions according to FANTOM CAT annotations using bedtools^103^. Magnification shows the distribution of LETR1 binding sites within promoter, exon, or intron of protein-coding genes (1,607 genes). LETR1 peaks are listed in Supplementary Table 9. **(d)** Venn diagrams showing the overlap between total genes expressed in LECs (TPM and CPM > 0.5) and identified LETR1-ChIRP genes, and the significant overlap between LETR1-ChIRP genes and differentially expressed genes after LETR1-ASOKD. **(e)** Circular plot showing genome-wide interactions of LETR1 near the 44 targets generated by Circos^104^. Scaled chromosomes with their respective cytobands are placed in circle. Major and minor ticks represent 50Mb and 10Mb, respectively. Orange and purple lines show interactions between LETR1 locus and its up- and downregulated targets, respectively. Green line highlights the genomic locus of LETR1. **(f)** Heat maps based on expression levels (CAGE-Seq, CPM) in control ASO and LETR1-ASOKD samples (up, 2 replicates), as well as in LECs and BECs (down, 2 replicates) of the 44 LETR1 targets (orange: upregulated, purple: downregulated). Color code for row Z-Score values on a scale from −1 to +1. Genes were ordered by log2FC values of ASOKD data and according to their differential expression between LECs and BECs.

To further elucidate the possible interactions between LETR1 and chromatin, we performed chromatin isolation by RNA purification followed by DNA sequencing (ChIRP-Seq)^54^. Cross-linked LECs were hybridized with two biotinylated probe sets (Odd and Even, internal control) tiling LETR1 (Supplementary Table 8). Probes targeting LacZ were used as an additional control. After pull-down, the percentage of retrieved RNA was assessed (Supplementary Figure 5b), and DNA was subjected to sequencing. Using a previously published analysis pipeline^54^, we found 2,258 binding sites of LETR1 to be at least 3-fold significantly enriched compared to input (P-value < 0.05; see Methods section), including a peak in the LETR1 exon one region as pull-down control (Supplementary Figure 5c and Supplementary Table 9).

To identify candidate genes directly regulated by LETR1, we first analyzed the genomic distribution of LETR1 binding sites. Out of 2,258 binding sites, 1,497 mapped within protein-coding genes (65.5%), with a large fraction residing in introns (1,010 peaks, 68.3%) (Figure 5c). Since only 19% of all annotated genes are categorized as protein-coding in the FANTOM CAT database^13^, these results suggested a preference of LETR1 to interact with regulatory regions near protein-coding genes (fold enrichment = 3.24, P-value < 0.05). Therefore, we focused on the identified 1,607 protein-coding genes displaying at least one LETR1 binding site within their promoters, exons, or introns. From these, 1,193 genes were expressed in LECs, and comparison with the 255 modulated genes upon LETR1-ASOKD showed a significant overlap of 44 genes (12 upregulated and 32 downregulated) (fold enrichment = 1.9, P-value < 0.05) (Figure 5d). Importantly, the vast majority of the 44 targets resided on different chromosomes, indicating a predominant *trans*-regulatory function of LETR1 (Figure 5e). These included important lymphatic-related genes such as KLF4, ROBO1, SEMA3A, SEMA3C, and CCBE1. Interestingly, 29 of the 44 targets showed a congruent higher expression in LECs for downregulated genes, or in BECs for upregulated genes, implicating these genes as potential downstream targets of LETR1 (Figure 5f).

### LETR1 regulates cell proliferation and cell migration through transcriptional regulation of KLF4 and SEMA3C

Among the 44 potential downstream targets of LETR1, KLF4 caught our attention as potential cell proliferation regulator given its well-established tumor suppressor role^55^ and the previously observed upregulation at the RNA level as well as increased TF binding activity upon LETR1 knockdown (Figure 3). Among cell migration regulatory molecules, we focused on one member of the semaphorin protein family, SEMA3C, that was previously shown to enhance migration in endothelial cells^56^.

To functionally characterize the relationship between LETR1 and KLF4 as well as SEMA3C, we performed the experimental strategies represented in Figure 6a. For KLF4, we analyzed the cell cycle progression of LECs after LETR1-ASOKD, followed by siRNA knockdown of KLF4. As expected, LETR1-ASOKD resulted in an upregulation of KLF4 as well as an increase of G0-arrested LECs. Consecutive knockdowns of LETR1 and KLF4 rescued this phenotype by significantly increasing the fraction of proliferating LECs. However, downregulation of KLF4 alone (by 70%) was not sufficient to consistently improve the proliferation activity of LECs (Figure 6b, c and Supplementary Figure 6a). For SEMA3C, we first ectopically overexpressed the SEMA3C protein in LECs, using a lentiviral vector (Supplementary Figure 6b). Subsequently, we analyzed the migratory behavior of infected LECs after LETR1-ASOKD. Again, LETR1-ASOKD alone caused the expected downregulation of SEMA3C as well as reduced migration in the vector-control cells. In contrast, overexpression of SEMA3C in conjunction with LETR1-ASOKD showed a significant recovery of migration capability, as compared to LETR1-ASOKD alone. SEMA3C overexpression alone did not affect cell migration of LECs (Figure 6d, e and Supplementary Figure 6c).

**Figure 6:**
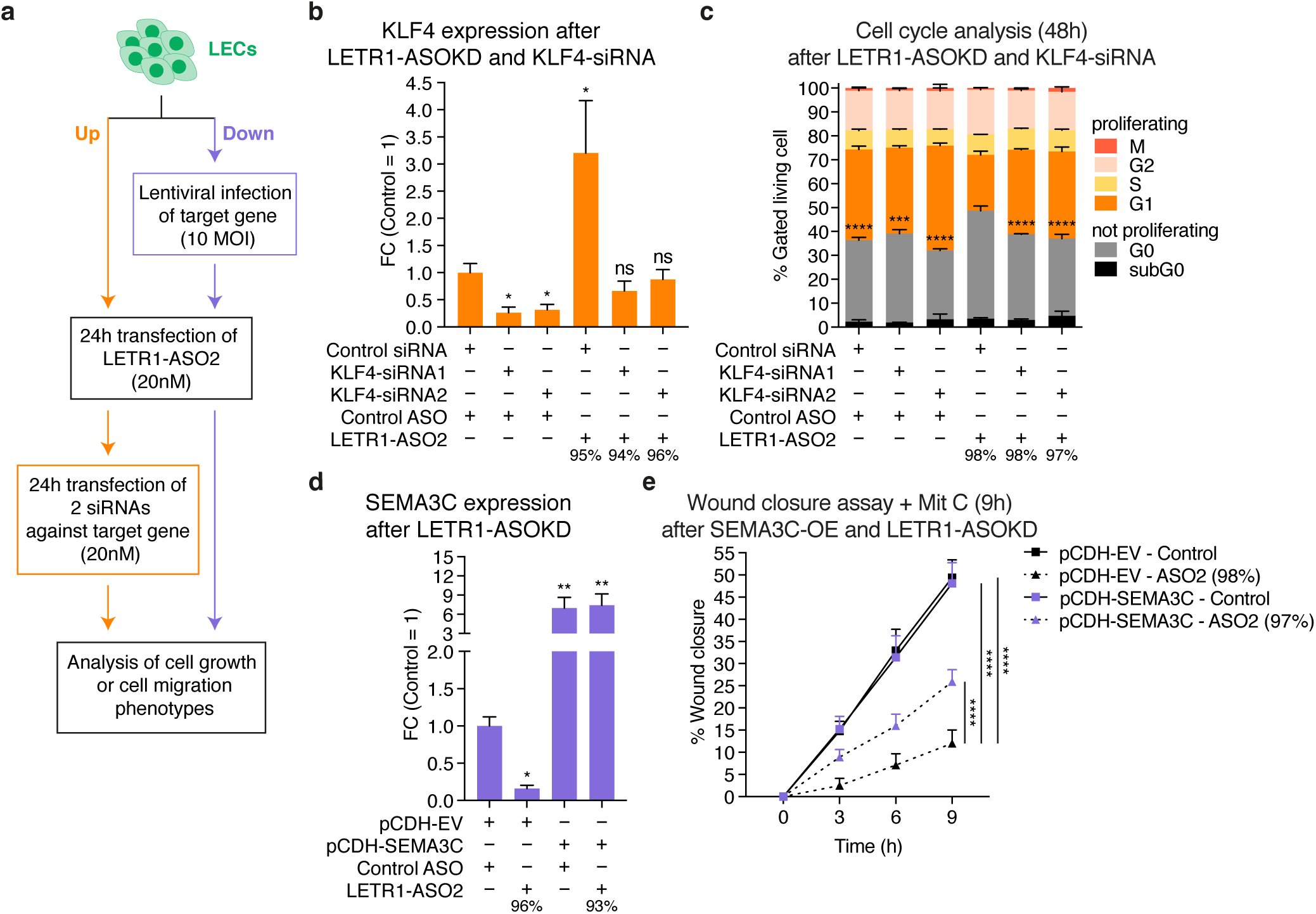
LETR1 regulates cell proliferation and cell migration through transcriptional regulation of KLF4 and SEMA3C. **(a)** Schematic representation of the experimental strategy to analyze the rescue of LETR1-ASOKD associated phenotypes with involved up- and down-regulated genes after combining LETR1 knockdown and gene targets dysregulation. **(b)** KLF4 expression levels after consecutive LETR1-ASOKD followed by siRNA-KD of KLF4 in neonatal LECs derived from 3 donors. **(c)** Quantification of the cell cycle progression analysis after 48h consecutive knockdown of LETR1 and KLF4. Statistical analysis was performed on G0 populations. **(d)** SEMA3C expression levels after LETR1-ASOKD in pCDH-EV and pCDH-SEMA3C infected neonatal LECs derived from 3 donors. **(e)** Quantification of wound closure assay (up to 9h) in neonatal LECs after the combination of SEMA3C overexpression and LETR1-ASOKD. Wound closure percentages were determined using TScratch^98^. Data are displayed as mean + SD (n = 3 in b-d; n = 10 in e). Percentages represent the knockdown efficiencies of LETR1 after the experiments. *P < 0.05, **P < 0.01, ***P < 0.001, ****P < 0.0001, ns not significant using RM one-way (for b, d), ordinary one-way (for c), and ordinary two-way (for e) ANOVA with Dunnett’s multiple comparisons test against control ASO – control siRNA (b), LETR1-ASO2 – control siRNA (c), pCDH-EV – control ASO (d), and pCDH-EV-ASO2 (e). All displayed *in vitro* assays were performed in neonatal LECs derived from the same donor.

### LETR1 interacts with several protein complexes to exert its regulatory function on gene expression

To identify proteins that are potential co-regulator of LETR1 target genes, we performed *in vitro* biotin-LETR1 pull-down assays^57^. Nuclear extracts of LECs were incubated with the biotinylated full-length LETR1 transcript and its antisense as negative control (Supplementary Figure 7a, b). After streptavidin bead separation, mass spectrometry was performed to identify possible interacting proteins. Initial analysis identified a total of 642 proteins. After filtering for proteins present in both replicates but absent in the antisense control, we found 59 proteins to interact with LETR1 (Supplementary Table 10). GO analysis for molecular functions and cellular compartments using g:ProfileR^36^ confirmed that the 59 identified proteins were significantly enriched for nuclear RNA-binding proteins (Supplementary Figure 7c, d). Protein-protein interaction analysis using STRING^58^ revealed that a large fraction of these 59 proteins were associated with RNA-processing functions, such as RNA splicing, RNA polyadenylation, and RNA nuclear transport. Furthermore, six proteins were associated with chromatin remodeling and three with nuclear organization, suggesting that LETR1 may operate at several levels to regulate gene expression (Figure 7a).

**Figure 7:**
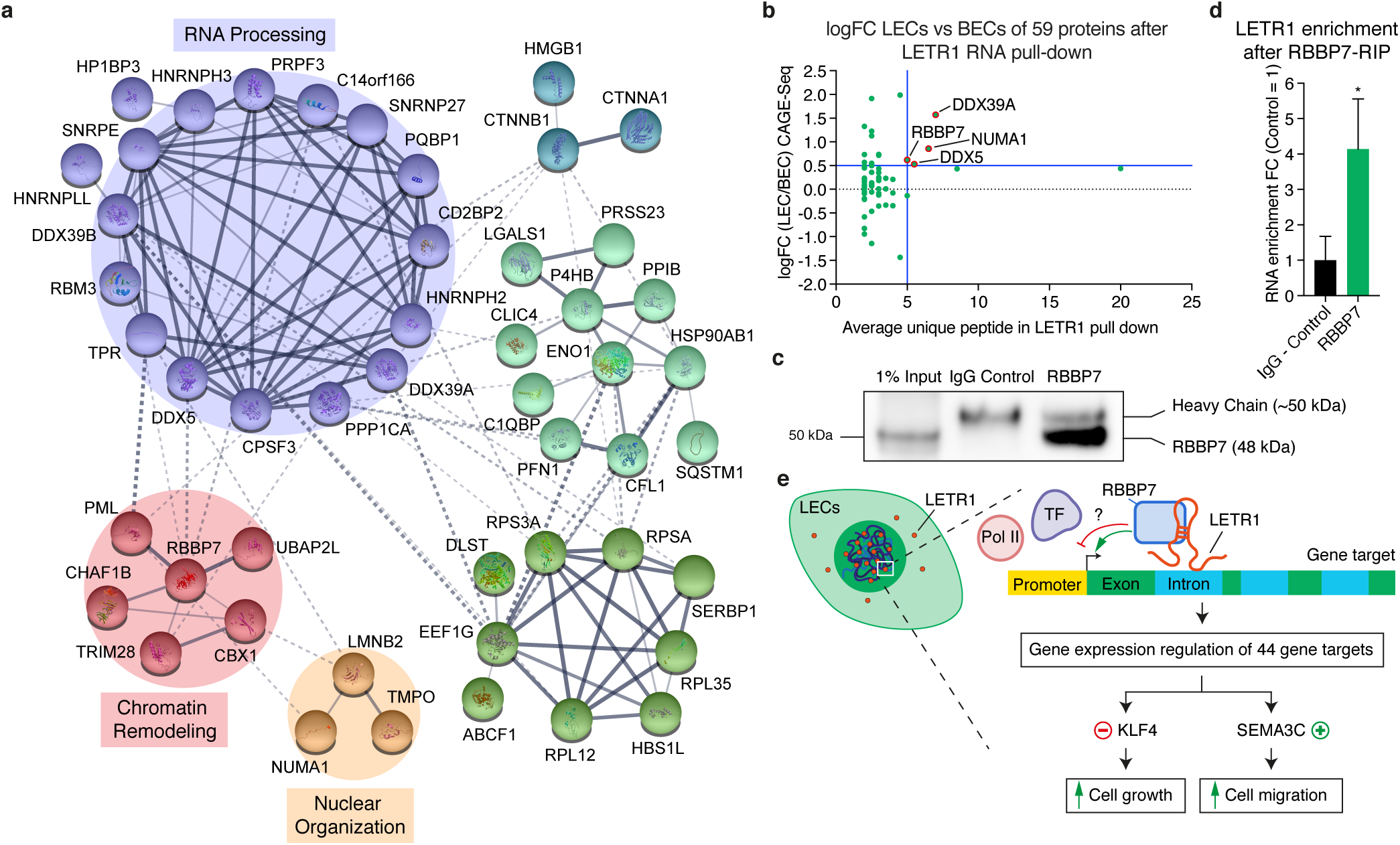
LETR1 interacts with several protein complexes to exert its regulatory function on gene expression. **(a)** Protein-protein interaction network using STRING^58^ of the 59 identified protein targets after *in vitro* biotin-RNA pull-down followed by mass spectrometry of LETR1 transcript and its antisense control in neonatal LECs derived from the same donor (n = 2). Proteins were clustered by Markov clustering (MCL) algorithm^107^ with an inflation parameter of 1.5. Circles highlight the most relevant clusters. Disconnected nodes were hidden to improve visualization. Lines indicate interactions within each complex. Thickness of the lines indicates the confidence of interaction from text mining, databases, experiments, and co-expression. Interacting proteins are listed in Supplementary Table 10. **(b)** Graph showing logFC LECs versus BECs against average unique peptide after LETR1 RNA pull-down. Blue lines show the logFC > 0.5 and unique peptide > 5 thresholds. **(c)** Representative western blot after RNA immunoprecipitation for RBBP7 followed by qPCR for LETR1 in neonatal LECs. To prevent masking from IgG heavy chain, a conformation-specific IgG secondary antibody was used. Uncropped western blot image is shown in Supplementary Figure 8. **(d)** LETR1 enrichment displayed as FC against IgG Control after RNA immunoprecipitation for RBBP7 in neonatal LECs derived from the same donor. LETR1 qPCR levels were normalized to the housekeeping gene GAPDH. Bars represent mean ± SD (n = 3). *P < 0.05 using unpaired Student’s t-test against IgG Control. **(e)** Model for the mode of action of LETR1 in the regulation of cell growth and cell migration in LECs through transcriptional regulation of 44 genes, including the validated KLF4 and SEMA3C.

To screen for protein candidates, we analyzed the RNA expression of the 59 proteins interacting with LETR1 in LECs versus BECs. Four proteins (DDX39A, NUMA1, RBBP7, and DDX5) had a logFC greater than 0.5 in LECs and a unique peptide detection greater than 5 (Figure 7b). Among these proteins, we identified the histone-binding protein RBBP7, which has previously been reported to be involved in the regulation of many cellular functions, including proliferation and migration^59,60^. Subsequent RNA immunoprecipitation assays in LECs validated the interaction between RBBP7 and LETR1, suggesting RBBP7 as a potential mediator of LETR1 gene regulatory functions (Figure 7c, d).

In summary, our multilayered mode of action analysis demonstrates that LETR1 is a nuclear lncRNA that interacts with essential epigenetic partners to regulate cell growth and cell migration of LECs by tuning the expression of distinct target genes, in particular KLF4 and SEMA3C (Figure 7e).

## Discussion

Precise regulation of proliferation, migration, and maintenance of cellular identity are not only essential to ensure proper development and integrity of the vascular systems, but also to guarantee that LECs and BECs are able to perform their necessary functions^5^. In this study, we characterized, for the first time, the global lineage-specific lncRNAome of LECs and BECs, and analyzed the transcriptional impacts after ASO-mediated knockdown of LEC- and BEC-specific lncRNAs followed by CAGE-Seq. Importantly, we identified LETR1 as a novel lymphatic-specific lncRNA that is essential in the regulation of lymphatic endothelial cell growth and migration.

By integrating RNA-Seq and CAGE-Seq transcriptome profiling, we showed that LECs and BECs express a specific cohort of lncRNAs, mainly residing near vascular-related protein-coding genes. These results are in accordance with the intriguing concept that cells might display a set of lncRNAs explicitly expressed to function in the fine-tuning of cell-type-specific gene expression programs^11^. Most notably, our selection strategy highlighted 2 LEC and 2 BEC lncRNAs that are robustly and differentially expressed in the respective endothelial lineage. These candidates therefore represent the first set of lineage-specific lymphatic and blood vascular endothelial cell lncRNA markers.

The nuclear localization of our lncRNA candidates coincides, to some extent, with previous findings, demonstrated by RNA in situ hybridization, that lncRNAs are commonly located in the nucleus^61^. To investigate the biological functions of these lncRNA candidates, we used the antisense oligonucleotide GapmeR knockdown approach, given its higher efficiency in targeting nuclear RNA transcripts over siRNA^38^. Additionally, our ASO design strategy proved to be very successful, resulting in an extremely high knockdown efficiency of all four targets. In the general experimental design, we have not taken into consideration the use of CRISPRi due to several practical limitations to study lncRNAs. For instance, as opposed to ASOKD, CRISPRi may also interfere with the transcription of overlapping or neighboring transcriptional units, and it is not able to distinguish the *cis* and *trans*-acting functions of lncRNAs^62,63^. However, we are aware that CRISPRi provides less pronounced off-target effects compared to other loss-of-function methods^64^.

Importantly, our study identified the first LEC lncRNA with specific biological functions. Through a multilayered analysis, including differential expression analysis, GO, GSEA, and MARA, we found that LETR1, originally annotated as LINC01197, is a critical gatekeeper of the global transcriptome of LECs by influencing complex TF regulatory networks regulating essential targets largely involved in the control of LEC growth and migration. These “molecular phenotypes” observed after LETR1 knockdown were confirmed *in vitro* by analyzing the cell growth profile, cell cycle progression, and wound closure ability of primary human LECs. As shown in a previous study^33^, we thus were able to distinguish LETR1 as a *bona fide* functional lncRNA in LECs by combining sequencing data analyses and cellular phenotype assays.

The nuclear localization of LETR1, as shown by subcellular fractionation and smRNA-FISH, hinted a chromatin-related function. Indeed, as confirmed by RNA-DNA interaction assay, we revealed that LETR1 interacts, predominantly *in trans*, with DNA regions near a subset of differentially expressed genes. In addition, RNA-protein interaction assays indicated a potential scaffold function of LETR1 in recruiting proteins involved in several levels of gene expression regulation, including chromatin organization. These results are in line with the general model in which lncRNAs are crucial for the assembly of unique protein complexes and for guiding them to specific target sites^65^. Specifically, we found that LETR1 interacts with RBBP7, a protein previously reported to be part of several multi-protein complexes that are involved in chromatin remodeling, histone post-transcriptional modification, and gene expression regulation^66-68^. Intriguingly, RBBP7 is a relevant constituent of the polycomb repressive complex 2 (PRC2) complex, which was also previously shown to interact with 20% of the lncRNAs in human cells^69^. It is conceivable that LETR1 might act as an epigenetic regulator to recruit or guide protein partners to influence the three-dimensional structure of the genome. In this setting, the differential CAGE-Seq peak intensities at the target TSSs after LETR1-ASOKD might provide a hint on the function of LETR1 in mediating the transcriptional machinery access at the site of transcription. In fact, significant correlations at TSS regions between RNA polymerase II occupancy and CAGE-Seq signal have been reported^70,71^.

We showed that LETR1 exerts its effects on LEC functions, at least in part, via the modulation of KLF4 and SEMA3C, since knockdown of KLF4 or overexpression of SEMA3C partially restored the cellular phenotypes observed upon LETR1-ASOKD. Previous studies have reported that KLF4 is a tumor-suppressor TF that, once upregulated, inhibits cell growth and induces cell cycle arrest^72-74^. A primary mechanism by which KLF4 regulates cell growth is via the induction of CDKN1A expression, a gene encoding a cyclin-dependent kinase (CDK) inhibitor^73,74^. Consistently, we found that CDKN1A was also upregulated upon knockdown of LETR1 (Supplementary Table 3), suggesting that suppression of the KLF4-CDK1NA axis through LETR1 is required for the maintenance of a proliferative state of LECs. Moreover, MARA analysis highlighted that the *trans*-acting activity of LETR1 has a general inhibitory effect on the binding activity of KLF4, suggesting an additional genome-wide interplay between KLF4 and LETR1 to modulate sensitive targets indispensable for LEC function.

A recent study reported that viral ectopic expression of FOXO1 in ductal pancreatic adenocarcinoma cells (PDAC) upregulates LINC01197 (LETR1), resulting in the inhibition of cell proliferation^75^. Interestingly, FOXO1 has previously been shown to promote LEC migration and to participate in the regulation of lymphatic development^76,77^. Moreover, in our sequencing data, FOXO1 was significantly more highly expressed in LECs than BECs.

SEMA3C belongs to the semaphorin class 3 guidance cue molecules, which mainly bind to a receptor complex composed of neuropilins (NRP1 or NRP2) and plexins (PLXNA1-A4 and PLXND1)^78^. Our findings that overexpression of SEMA3C partially rescued the LETR1 inhibition of LEC migration are congruent with previous reports showing that SEMA3C has pro-migratory activities in several cell types^79-81^, including endothelial cells^82^. In support of this claim, our sequencing data revealed that LECs expressed the two neuropilins, NRP1 (at a low level) and NRP2, as well as the plexins A1-A4 and D1. Additionally, knockdown of LETR1 also significantly reduced the expression of plexin A4 (Supplementary Table 3), pinpointing LETR1 as an intermediate player in semaphorin signaling.

While we initially identified LETR1 as a LEC-specific lncRNA by sequencing of cultured human LECs, smRNA-FISH and FACS validated its lymphatic specificity in human skin *in situ*. Future studies are needed to investigate its expression pattern and mechanistic role in pathological conditions associated with impaired lymphatic function (e.g., lymphedema), or active lymphangiogenesis (e.g., tumor metastasis and wound healing).

It is of interest that the knockdown of three other lncRNA candidates showed merely a minor or no impact on the transcriptome of LECs and BECs after ASO-mediated knockdown. Likely, these could be due to four potential reasons. First, they may have alternative functions unrelated to transcriptional regulation, such as ribozymes or riboswitches^83^ and translation initiation regulators^84^. Second, the act of transcription, rather than the lncRNA product of this transcription, may be functional by having, for instance, an enhancer-like function^85^. Third, they may function as molecular signals at a specific time and place in response, for example, to unique stimuli^86^. Finally, although clearly differentially expressed, all three lncRNA candidates might not be functional and might just be part of transcriptional noise^87^. Therefore, future research is needed to elucidate the biological role and function of these lncRNAs in LECs or BECs.

Taken together, our study enumerates the collection of lncRNAs explicitly expressed in LECs and BECs and highlights, through the functional characterization of LETR1, the importance of those lncRNAs in the regulation of lineage-specific endothelial cell functions.

## Materials and Methods

### Isolation of adult primary skin LECs and BECs from biopsies

LECs and BECs were obtained from the abdominal or breast skin of healthy adult subjects admitted for plastic surgery at the University Hospital Zurich. Written informed consent was obtained from each donor/tissue collection, as approved by the Ethics Committee of the Kanton Zurich (2017-00687). Skin samples were washed in hank’s balanced salt Solution (HBSS) supplemented with 5% fetal bovine serum (FBS, Gibco), 2% antibiotic and antimycotic solution (AA, Gibco) and 20mM HEPES (Gibco), and subsequently incubated in 0.25% trypsin (Sigma) diluted in DPBS (Gibco) with the dermal side facing downwards overnight at 4°C. Trypsin digestion was stopped by washing the tissues with RPMI basal medium supplemented with 10% FBS, 2% AA, and 20mM HEPES. After removal of the epidermal sheets, the dermis was finely minced and enzymatically digested (RPMI basal medium, 1000U/mL collagenase type 1 (Worthington), 40*µ*g/mL DNase I (Roche)) for 1h at 37°C under constant agitation. Digested tissues were then filtered through a 100*µ*m cell strainer (Falcon), washed with RPMI basal medium, and centrifuged at 1,500rpm for 6min at 4°C. Cells were seeded into fibronectin (Roche) coated plates and were cultured in EGM-2-MV complete medium (Lonza). After 7-10 days, cells were trypsinized, and endothelial cells were selected based on CD31 positivity with Dynabeads CD31 endothelial cell magnetic beads (Thermo Fisher Scientific) and cultured until confluency. Endothelial cells were detached, washed with FACS buffer (DPBS with 2% FBS and 1mM EDTA), and stained with Alexa647-conjugated mouse anti-human podoplanin antibody (1:70, clone 18H5, Novus Biologicals) and PE-conjugated mouse anti-human CD31 antibody (1:20, clone WM59, BD Pharmingen) in FACS buffer for 30min at 4°C. After a wash with FACS buffer, endothelial cells were finally sorted on a FACSAria II (BD Biosciences) with a 70*µ*m nozzle, using FACSDiva software. LECs were defined as CD31- and podoplanin-positive cells, whereas BECs were defined as CD31-positive and podoplanin-negative cells.

### Cell culture

Primary human dermal LECs and BECs were isolated from neonatal human foreskin as described previously^88^ and cultured in endothelial basal medium (EBM, Lonza) supplemented with 20% FBS, 100U/mL penicillin and 100*µ*g/mL streptomycin (Pen-Strep, Gibco), 2mM L-glutamine (Gibco), 10*µ*g/mL hydrocortisone (Sigma) on sterile dishes/plates (TPP) pre-coated with 50*µ*g/mL purecol type I bovine collagen solution (Advanced BioMatrix) in DPBS at 37°C in a 5% CO_2_ incubator. LECs were additionally cultured in the presence of 25*µ*g/mL cAMP (Sigma); BECs in the presence of endothelial cell growth supplement ECGS/H (PromoCell). Cells were used at passage 6-7 for RNA-Seq and CAGE-Seq experiments, and at passage 8-9 in *in vitro* and biochemistry experiments. Primary human dermal LECs and BECs were isolated from adult human skin as described above and cultured in EGM-2 complete medium in vessels coated with fibronectin diluted in DPBS and cells were cultured at 37°C in a 5% CO_2_ incubator. Cells were used between passage 2 and 6. HEK293T cells were cultured in DMEM with glutamax (Gibco) supplemented with 10% FBS (Gibco) and Pen-Strep at 37°C in a 5% CO_2_ incubator. All cells were routinely tested for mycoplasma contamination using the MycoScope PCR Mycoplasma Detection kit (Genlantis).

### RNA isolation, reverse transcription, and qPCR

If not differently specified, total RNA was isolated using the NucleoSpin RNA kit (Machery Nagel), according to the manufacturer’s instructions and quantified by NanoDrop ND-1000 (Witec AG). Equal amounts of total RNA were reverse transcribed using the High Capacity cDNA Reverse Transcription Kit (Applied Biosystems), according to the manufacturer’s instructions. 10ng cDNA per reaction was then subjected to qPCR using PowerUp SYBR Green Master Mix (Applied Biosystems) on a QuantStudio 7 Flex Real-Time PCR system (Applied Biosystems). For qPCR analysis, cycle threshold (Ct) values were normalized to the housekeeping gene RPLP0 or GAPDH. Relative expression was calculated according to the comparative Ct method. Primers are listed in the Supplementary Table 11.

### Western blot analysis

To perform western blot analysis, the protein concentration of lysates was first determined using the Microplate BCA protein assay kit – reducing agent compatible (Thermo Fisher Scientific), according to the manufacturer’s instruction. To 5-30*µ*g of total protein, SDS sample buffer and reducing agent (Thermo Fisher Scientific) were added to a 1x final concentration. Then, samples were heated for 5min at 70°C and size-separated by electrophoresis using 1.5mm 4-12% NuPAGE Bis-Tris protein gels and 1x NuPAGE MES SDS running buffer (Thermo Fisher Scientific) for 35-50min at 200V. Proteins were transferred to a nitrocellulose membrane (Merck Millipore) for 1h at 20V. Protein loading was checked by staining membranes with Ponceau staining solution (Sigma) for 2min at room temperature. Membranes were blocked with 5% milk powder in TBST (50mM Trizma Base, 150mM NaCl, 0.1% Tween 20, pH 8.4) for 1.5h at room temperature. Membranes were then stained overnight at 4°C with primary antibodies (see below) diluted in TBST. Blots were washed three times with TBST for 15min at room temperature and subsequently incubated for 2h at room temperature with HRP-conjugated secondary antibodies (goat anti-rabbit, Dako; rabbit anti-goat, R&D Systems) at a dilution of 1:5000 in TBST. After washing five times with TBST for 15min at room temperature, blots were developed with clarity western ECL substrate (Bio-Rad) and imaged on a ChemiDoc imaging system (Bio-Rad). All antibodies are listed in the Supplementary Table 11.

### Lentivirus production

For production of lentiviruses, 2.5×10^6^ HEK293T cells were seeded into 10cm dishes and cultured overnight. One hour before transfection, the medium was replaced with an antibiotic-free medium containing 25*µ*M chloroquine (Sigma). The transfection mixture was subsequently prepared as follows. In a first tube, 1.3pmol psPAX2 (12260, Addgene), 0.72pmol pMD2.G (12259, Addgene), and 1.64pmol of target vector were mixed in 500*µ*L Opti-MEM (Gibco). In another tube, polyethylenimine (PEI, Sigma) was added to 500*µ*L Opti-MEM in a 1:3 ratio to total DNA content. PEI-containing Opti-MEM was transferred dropwise to the plasmid-containing Opti-MEM, and the mixture was incubated for 20min at room temperature. Finally, the transfection mixture was transferred dropwise to the HEK293T cells. 24h post-transfection, the medium was changed with 8mL complete medium containing 10*µ*M forskolin (Sigma). Lentiviruses were harvested after 48h, filtered with a 0.45*µ*m PES filter (TPP), and stored at −80°C. The titer of each virus was determined using the Lenti-X Go-Stix Plus kit (Takara Bio), according to the manufacturer’s instruction.

### RNA-Seq and CAGE-Seq of primary LECs and BECs

2.5×10^5^ LECs and BECs were seeded in duplicates into 10cm dishes and cultured for three days until 70% confluence was reached. At this point, 8mL EBM consensus medium (EBM supplemented with 20% FBS, L-glutamine, and Pen-Strep) was added to both cell types. After 48h, total RNA was harvested and isolated using RNeasy mini kit (Qiagen). DNA digestion was performed using the RNase-Free DNase set (Qiagen). The identity of BECs and LECs was checked by qPCR (Supplementary Figure 1a, b). Primers are listed in the Supplementary Table 11. LEC and BEC total RNA were then subjected to ribosomal-RNA depleted RNA sequencing (RNA-Seq) and nAnT-iCAGE sequencing (CAGE-Seq) protocols as previously described^31^.

### Differential gene expression analysis of CAGE-Seq and RNA-Seq data

For both RNA-Seq and CAGE-Seq data, read alignment was performed, and expression tables were generated as described previously^33^. Next, we performed differential expression (DE) analysis of RNA-Seq and CAGE-Seq of LECs against BECs, LECs against DFs, and BECs against DFs using EdgeR (ver. 3.12.1)^34,89,90^. LEC-associated genes: false discovery rate (FDR) < 0.01 and log2 fold change (FC) LECs/BECs > 1 and log2FC LECs/DFs > 1. BEC-associated genes: FDR < 0.01 and log2FC LECs/BECs < −1 and log2FC BECs/DFs > 1. LEC- and BEC-associated lncRNAs were selected according to their annotation as “lncRNA” in the FANTOM CAT database^13^. LEC and BEC core lncRNAs were then defined as the overlap between RNA-Seq and CAGE-Seq analyses.

### Genomic classification and flanking gene analyses of LEC and BEC core lncRNAs

To analyze the genomic classification of LEC and BEC core lncRNAs, we first determined which lncRNAs were broadly expressed in either BECs or LECs by applying an expression level cut-off of 0.5 on both TPM (RNA-Seq) and CPM (CAGE-Seq). Next, LEC/BEC core lncRNAs and broadly expressed lncRNAs were classified according to the “CAT gene class” category of the FANTOM CAT database^13^. Finally, enrichment analysis of LEC/BEC intergenic lncRNAs versus broadly expressed intergenic lncRNAs was performed using SuperExactTest (ver. 1.0.0)^91^. As background, total annotated lncRNA in FANTOM CAT database^13^ (n = 90,166) were used.

To determine flanking protein-coding genes, we uploaded lists containing transcriptional start sites (TSSs) of LEC/BEC core lncRNAs to the GREAT webtool (http://great.stanford.edu/public/html/, ver. 4.0.4)^35^. As association rule, we used the “two nearest genes” option with a maximal extension from the lncRNA TSS of 10 Mb. Gene Ontology (GO) analysis was then performed on determined protein-coding genes, using g:Profiler (ver 0.6.7)^36^. Genes with TPM (RNA-Seq) and CPM (CAGE-Seq) < 0.5 were used as background. The gprofiler database Ensembl 90, Ensembl Genomes 37 (rev 1741, build date 2017-10-19) were used. Only GO terms with P-value < 0.05 were used for further analysis.

### Antisense oligonucleotide design and efficiency test

Five locked nucleic acids (LNA) phosphorothioate GapmeRs per lncRNA target were designed as described previously^33^. After determining the TSS of each target by evaluating their CAGE-Seq signals, ASOs were placed in the first intronic region downstream of the identified TSS (Supplementary Figure 2). ASO sequences are listed in Supplementary Table 2. To test the knockdown efficiency of each ASO, 35,000 LECs per well were seeded into a 12-well plate and cultured overnight. LECs were then transfected with 20nM of scramble control ASO or five ASOs (GeneDesign) targeting AL583785.1, LETR1, LINC00973, or LINC01013, and 1*µ*L Lipofectamine RNAiMAX (Thermo Fisher Scientific) previously mixed in 100*µ*L Opti-MEM according to the manufacturer’s instructions. Knockdown efficiency for each ASO was checked by qPCR, as described above. Only the three most potent ASOs per target were used in the ASOKD screen, followed by CAGE-Seq (Supplementary Figure 2).

### ASOKD-screen in LECs and BECs followed by CAGE-Seq

7×10^5^ LECs and 6×10^5^ BECs were seeded into 10cm dishes and cultured overnight. Subsequently, the growth medium of both cell types was replaced by 8mL EBM consensus medium. LECs and BECs were then cultured for additional 24h. To transfect LECs and BECs, 20nM of each ASO (3 ASOs per target plus scramble control ASO) and 16*µ*L Lipofectamine RNAiMAX were mixed in 1.6mL Opti-MEM according to the manufacturer’s instructions. After 5min incubation at RT, the Lipofectamine-ASO mixture was added dropwise to the cells. LECs and BECs were harvested after 48h post-transfection. For these samples, total RNA was isolated using the RNeasy mini kit. DNA digestion was performed using the RNase-Free DNase set. Knockdown efficiency for each ASO was checked by qPCR, as described above. Samples with at least 50% knockdown efficiency were subjected to CAGE-Seq. Knockdown efficiency was also confirmed after CAGE-Seq (Supplementary Figure 3a, b).

### Differential gene expression analysis of ASOKD data

All ASOs for each targeted lncRNA were compared against scramble control ASO (NC_A) libraries from corresponding cell types. Genes with expression >= 5 CPM in at least two CAGE libraries (targeted lncRNA ASOs + scramble control ASO (NC_A) CAGE libraries) were tested for differential expression (DE). A generalized linear model (GLM) was implemented for each targeted lncRNA to perform DE analysis using EdgeR (ver. 3.12.1)^34,89,90^. Genes with |log2FC| > 0.5 and FDR < 0.05 were defined as differentially expressed genes and used for the downstream analysis.

### Gene ontology (GO) analysis of ASOKD data

GO analysis was performed separately on upregulated and downregulated genes, using g:Profiler (ver 0.6.7)^36^, as described above. Genes with expression >= 5 CPM in at least two CAGE libraries were used as background. All the significant GO terms (P-value < 0.05) were used for further analysis and are listed in Supplementary Table 4.

### Gene set enrichment analysis (GSEA) of ASOKD data

GSEA was performed individually for each targeted lncRNA using tool xtools.gsea.Gsea from javascript gsea2-2.2.4.jar^45,92^. All ASOs for each targeted lncRNA were compared against scramble control ASO (NC_A) libraries from corresponding cell types. Genes with expression >= 5 CPM in at least two CAGE libraries were included in the input table for the analysis. Gene sets for GO (Biological Process, Molecular Function, Cellular Component), Hallmark, KEGG, Reactome, BioCarta, and Canonical pathways from MSigDB (ver. 6.0) were used for the analysis. The parameters used for each run were: -norm meandiv -nperm 500 -permute gene_set -rnd_type no_balance - scoring_scheme weighted -metric Signal2Noise -rnd_seed timestamp -set_max 1000 -set_min 5. Enriched GO biological processes were selected and organized in a network using Cytoscape (ver. 3.6.1) and the plugin Enrichment Map^93^. Gene-set filtering was set as following: FDR q-value cutoff < 0.05 and P-value cut off < 0.001. Gene-set similarity cutoff was set as < 0.5 with an Overlap Metric. Genes were filtered by expression. Terms were then organized manually according to their biological meaning using the Cytoscape plugin Wordcloud^93^. Significant GO biological processes after GSEA are listed in Supplementary Table 5.

### Motif activity response analysis (MARA) of ASOKD data

MARA was performed for BEC and LEC separately using promoter expression for all the knockdown (KD) and scramble control ASO (NC_A) libraries. All promoters with expression >=1TPM at least in 5 CAGE libraries were used for the analysis. MARA was performed as described previously^33^ for the motifs in SwissRegulon (released on 13 July 2015)^94^. Student’s t-test was performed to identify differentially active motifs due to the lncRNA-ASOKD. Motifs with significant motif activity differences (P-value < 0.05) compared to the controls (NC_A) were used for downstream analysis. Significant TF motifs after MARA are listed in Supplementary Table 6.

### Cloning sgRNA targeting LETR1 and establishment of dCas9 expressing LECs

sgRNAs targeting LETR1 were designed using the online CRISPR design tool from the Zhang lab, MIT (http://crispr.mit.edu/). 250bp upstream of the highest CAGE-Seq peak were used as the design region. We then selected then 3 sgRNAs to be cloned into lentiGuide-Puro (52963, Addgene) as previously described^95^. Briefly, each pair of oligos was first annealed and phosphorylated using T4 PNK (New England BioLabs) using the following program: 30min at 37°C, 5min at 95°C, and then ramped down to 25°C at 5°C/min. Annealed oligos were then diluted 1:200. LentiGuide-Puro vector was digested with BsmBI (New England BioLabs) and dephosphorylated using rSAP (New England BioLabs) for 4h at 37°C. After gel purification, LentiGuide-Puro linearized vector and annealed oligos were ligated using Quick Ligase (New England BioLabs) for 20min at room temperature. Ligated vectors were then transformed into one shot TOP10 chemically competent cells (Thermo Fisher Scientific), according to the manufacturer’s instruction. Plasmids were isolated using the Nucleospin Plasmid kit (Machery Nagel), as described in the manufacturer’s protocol. Sequences of inserted sgRNAs were checked by Sanger sequencing (Microsynth). Sequences of sgRNAs are listed in the Supplementary Table 11. Lentiviruses containing pHAGE EF1a dCas9-KRAB (50919, Addgene, with custom blasticidin cassette), scramble control sgRNA, or each of the sgRNA targeting LETR1 were produced as described above.

To establish dCas9-overexpressing LECs, 1.2×10^5^ LECs were seeded into pre-coated 6-well plates (TPP) and infected with medium containing dCas9-KRAB lentiviruses diluted at a 10 multiplicity of infection (MOI) and 5*µ*g/mL polybrene (hexadimethrine bromide, Sigma). Plates were then sealed with parafilm and centrifuged at 1,250rpm for 1.5h at room temperature. The next day, the medium was changed, and positively infected cells were selected with 10μg/mL blasticidin (InvivoGen). Once confluent, at least 5×10^5^ dCas9-KRAB-expressing LECs were split into pre-coated 10cm dishes and cultured under antibiotic selection until confluency. After checking RNA and protein levels of dCas9-KRAB as described previously^95^, LECs were then used in the cell growth profile experiment.

### Cell growth profiling after LETR1-ASOKD and -CRISPRi

For cell growth profiling after ASOKD, 3,000 LECs per well were seeded into a 96-well plate and cultured overnight. LECs were then transfected with 20nM of scramble control ASO or three ASOs targeting LETR1 (Exiqon) and 0.2*µ*L Lipofectamine RNAiMAX previously mixed in 20*µ*L Opti-MEM according to the manufacturer’s instructions.

For cell growth profiling after CRISPRi, 3,000 dCas9-expressing LECs per well were seeded into a 96-well plate and grown overnight. LECs were then infected with 50 MOI of lentiviruses containing vectors expressing scramble control sgRNA or three sgRNAs targeting LETR1 diluted in complete growth EBM medium supplemented with 5*µ*g/mL polybrene. After 24h, the virus-containing medium was changed.

In both experiments, LECs were continuously imaged every 3h over three days with 4 fields per well using the IncuCyte ZOOM live-cell imaging system (Essen Bioscience). Confluence in each well was determined using IncuCyte ZOOM software. The normalized growth rate was calculated as the slope of linear regression and normalized to control. To check knockdown efficiency, LECs were harvested after 72h post-transfection. Total RNA was isolated using the RNeasy mini kit. qPCR was performed using One Step SYBR PrimeScript RT-PCR kit (Takara Bio) on a 7900HT real-time system (Applied Biosystems).

### Cell cycle analysis after LETR1-ASOKD

Cell cycle analysis was adapted from^96^. 5×10^5^ LECs were seeded into 10cm dishes and cultured overnight in starvation medium (EBM supplemented with 1% FBS). The next day, LECs were transfected with scramble ASO or three ASOs targeting LETR1, as described above. As additional controls, LECs were incubated in the starvation medium as non-proliferative control or treated with 100ng/mL nocodazole (Sigma) as mitosis control. 24h post-transfection, LECs were detached and collected. Floating LECs were also collected in the same tube. 2×10^5^ LECs per replicate were then transferred into a 96 U-bottom plate (Greiner bio-one). Aliquots of approx. 1×10^5^ LECs were lysed and subjected to RNA isolation, reverse transcription, and qPCR, as described above. After a DPBS wash, LECs were stained with Zombie NIR (BioLegend) diluted 1:500 in DPBS for 15min at room temperature in the dark. Subsequently, LECs were washed with DPBS and fixed in 70% ethanol overnight at −20°C. After permeabilization, LECs were washed twice in DPBS and stained with mouse anti-human Ki-67 antibody (Dako) diluted 1:800 in FACS buffer (DPBS with 1mM EDTA and 2% FBS) for 30min at room temperature in the dark. As a negative control, one sample was stained with mouse IgG isotype control (clone 11711, R&D Systems) diluted 1:5 in FACS buffer. After two washes in DPBS, LECs were then stained with donkey Alexa488-conjugated anti-mouse secondary antibody (Thermo Fisher Scientific) diluted 1:500 in FACS buffer for 30min at room temperature in the dark. Next, LECs were washed once in DPBS and incubated with 20*µ*g/mL RNase A (Machery-Nagel) diluted in DPBS for 60min at room temperature. Finally, LECs were incubated with 10*µ*g/mL propidium-Iodide diluted in staining buffer (100mM Tris, 150mM NaCl, 1mM CaCl_2_, 0.5mM MgCl_2_, and 0.1% Nonidet P-40) for 20min at RT. Flow cytometry was performed with a CytoFlex S instrument (Beckman Coulter). Data were analyzed with FlowJo (ver. 10.lr3). Gating was done using live/dead to gate living cells, isotype control to gate proliferative/non-proliferative cells, starvation medium-treated sample to gate G0 population, and nocodazole-treated sample to gate mitotic cells.

### Apoptosis assay after LETR1-ASOKD

3,000 LECs per well were seeded into a 96-well plate and cultured overnight. LECs were then transfected with 20nM of scramble control ASO or three ASOs targeting LETR1, as described above. As a positive control, additional cells were treated for 2h with 4*µ*M staurosporine (Sigma). 48h post-transfection, LECs were fixed in 3.7% PFA (AppliChem) in DPBS for 20min at room temperature. After three washes with DPBS, LECs were blocked in blocking buffer (DPBS with 5% donkey serum (Sigma) and 0.3% triton X-100) for 1h at room temperature. LECs were then stained with rabbit anti-cleaved caspase 3 (Asp175) antibody (Cell Signaling) diluted 1:400 in antibody buffer (DPBS with 1% BSA and 0.3% triton X-100) overnight at 4°C. The next day, LECs were washed three times for 5min in DPBS and subsequently stained with donkey Alexa488-conjugated anti-rabbit secondary antibody (Thermo Fisher Scientific) diluted 1:1000, and Hoechst dye (Thermo Fisher Scientific) diluted 1:2000 in antibody buffer for 1h at room temperature in the dark. After three washes for 5min in DPBS, the plate was imaged using a fluorescence microscope (Zeiss Axiovert 200M), and four images at a 5X magnification were taken for each well. Percentage of cleaved caspase 3-positive cells was determined using a self-built macro developed with ImageJ (ver. 2.0.0-rc-69/1.52i)^97^. To check knockdown efficiency, an extra plate was lysed at the end of the experiment and subjected to qPCR, as described above.

### Wound closure assay after LETR1-ASOKD

2.5×10^5^ LECs were seeded into 6cm dishes and cultured for at least 6h. LECs were then transfected with 20nM of scramble control ASO or three ASOs targeting LETR1 and 8*µ*L Lipofectamine RNAiMAX previously mixed in 800*µ*L Opti-MEM according to the manufacturer’s instructions. 24h post-transfection, LECs were detached, and 15,000 were seeded into 96-well plates and cultured overnight. The next day, LECs were incubated for 2h with 2*µ*g/mL mitomycin C (Sigma) in complete EBM medium. After 2h incubation with proliferation inhibitor, LECs were scratched using a wounding pin replicator (V&P Scientific), according to the manufacturer’s instructions, complete EBM medium was replaced, and LECs were incubated for 9h. Images of the scratched areas were taken at 5x magnification and at time points 0h, 3h, 6h, and 9h using a bright field microscope (Zeiss Axiovert 200M). Images were analyzed using TScratch (ver. 1.0)^98^. To check knockdown efficiency, LECs were harvested after 9h, and total RNA, cDNA synthesis, and qPCR were performed as described above.

### Trans-well migration assay after LETR1-ASOKD

2.5×10^5^ LECs were seeded into 6cm dishes and cultured for at least 6h. Transfection of scramble control ASO or three ASOs targeting LETR1 was performed as described above. 24h post-transfection, 50,000 LECs were seeded in starvation medium into a trans-well (24-well plate, 6.5mm insert, 8*µ*m polycarbonate membrane, Costar) previously coated with collagen-I on both sides of the membrane. To check knockdown efficiency, an aliquot of each condition was lysed and subjected to qPCR, as described above. After 4h incubation at 37°C, LECs were fixed with 3.7% PFA in DPBS for 10min. After a DPBS wash, nuclei were stained with Hoechst dye diluted 1:1000 in DPBS for 10min at room temperature in the dark. LECs on the upper side of the membrane were then removed with cotton swabs, and the membrane was thoroughly washed with DPBS. Finally, membranes were cut and mounted on microscope slides with mowiol 4-88 (Calbiochem). For each membrane, four images were taken at 5x magnification using a fluorescence microscope (Zeiss Axioskop 2 mot plus). Cell migration was quantified by counting the nuclei per image field using a self-built macro developed with ImageJ (ver. 2.0.0-rc-69/1.52i)^97^.

### Subcellular fractionation followed by qPCR

Fractionation of LECs or BECs was adapted from^99^. After trypsinization, 1×10^6^ LECs or BECs were collected in 15mL Falcon tubes and washed once with DPBS. LECs or BECs were then resuspended in 1mL cold cell disruption buffer (10mM KCl, 1.5mM MgCl_2_, 20mM Tris-Cl pH 7.5, 1mM DTT) and incubated for 10min on ice. At this point, LECs or BECs were transferred into a 7mL Dounce homogenizer (Kimble) and homogenized with pestle type B for 25-30 times until free nuclei were observed under the microscope (Zeiss Axiovert 200M). LEC and BEC nuclei were subsequently transferred to a fresh tube, and triton X-100 was added to a final concentration of 0.1%. After mixing four times by inverting the tube, LEC and BEC nuclei were pelleted, and the supernatant was recovered as cytoplasmic fraction. To isolate nuclear RNA, the nuclei pellet was lyzed in 1mL GENEzol reagent (Geneaid Biotech). After 5min incubation at room temperature, 200*µ*L chloroform were added to the homogenized nuclear fraction. After vortexing for 10s, the nuclear fraction was spun down at 13,000rpm for 15min at 4°C. The upper aqueous phase was transferred into a new tube. For cytoplasmic RNA, on the other hand, two volumes of phenol:chlorofrom:isoamyl alcohol mixture (PCA, 25:24:1, Sigma) were added to the cytoplasmic fraction and vortexed for 1min. After spinning for 5min at 4,000rpm, the upper aqueous phase was transferred into a tube. One volume of isopropanol was added to both the nuclear and cytoplasmic aqueous phases and mixed by inverting the tubes several times. After 10min incubation at room temperature, the tubes were centrifuged at 13,000rpm for 10min at 4°C. RNA pellets were subsequently washed with 75% ethanol and dried for 10min at room temperature. Dried RNA pellets were resuspended in RNase-free water and incubated for 15min at 58°C on a heating block. Finally, both samples were subjected to cDNA synthesis, and qPCR was performed as described above.

### Identification of transcripts variants of LETR1 in LECs

The transcriptional start site (TSS) of LETR1 was determined by examining the CAGE-Seq signal (Supplementary Figure 2). Transcripts rising from the identified TSS were determined using the SMARTer Rapid amplification of cDNA ends (RACE) 5’/3’ kit (Takara Bio) in accordance with the manufacturer’s instructions for 3’ RACE. 100ng cDNA synthesized from total LEC RNA was used in 3’ RACE reactions. The primer used in the 3’ RACE assay was designed using CLC Genomic Workbench (ver. 10.1.1, Qiagen) near the highest CAGE-Seq signal in the determined TSS region. The primer sequence was 5’-GATTACGCCAAGCTTTTGTGAGCCACTGCGTTCT-3’, and the annealing temperature was 62°C. Polyadenylation of LETR1 transcripts was determined using qPCR and comparing the expression values between cDNA synthesis with random and oligodT_20_ primers. Expression levels of the identified primers were determined using qPCR as described above. Primers to amplify LETR1 transcripts are listed in the Supplementary Table 11. Sequences of LETR1 transcript variants are listed in Supplementary Table 7.

### Simultaneous RNA fluorescence in situ hybridization (FISH) and immunostaining in cultured LECs

5,000 LECs per well were seeded into a 96-well glass-bottom imaging plate (Greiner bio-one). Once confluence was reached, smRNA-FISH was performed using the viewRNA Cell Plus Assay kit (Thermo Fisher Scientific), according to the manufacturer’s instructions. LECs were stained with probes designed to target human LETR1 (Type 1 Probe, VA1-3018146, Thermo Fisher Scientific), human Malat1 (positive control, Type 1 Probe, VA1-11317, Thermo Fisher Scientific), and bacterial DapB (negative control, Type 1 Probe, VF1-11712, Thermo Fisher Scientific). LECs were additionally co-stained with mouse anti-human CD31 antibody (clone JC70A, Dako) at a dilution of 1:50, followed by donkey anti-mouse Alexa-Fluor 488 secondary antibody (Thermo Fisher Scientific) at a dilution of 1:200. Z-stacks of fluorescence images spanning over the entire cell monolayer were acquired using an inverted confocal microscope (Zeiss LSM 780). A self-built macro developed with ImageJ (ver. 2.0.0-rc-69/1.52i)^97^ was used to quantify the nuclear versus cytoplasmic localization of LETR1 by applying a max intensity projection. After determining the nuclear surface using the Hoechst dye signal, plugin “analyze particles” was used to count spots present either in the nuclear or cytoplasmic area.

### Simultaneous smRNA-FISH and immunostaining in human skin tissues

Normal human skin samples were obtained from plastic surgery. Immediately after dissection, samples were fixed in fresh 10% neutral buffered formalin (NBF) for 24h at room temperature. Fixed samples were then dehydrated using a standard ethanol series followed by xylene, and were embedded in paraffin. Using a microtome, 5*µ*m skin sections were cut and mounted on superfrost plus slides (Thermo Fisher Scientific). Slides were dried overnight at room temperature. Simultaneous smRNA-FISH and immunostaining was performed using the RNAScope Multiplex Fluorescent Reagent kit v2 (Advanced Cell Diagnostics), according to the manufacturer’s instructions and technical note 323100-TN. Briefly, slides were backed for 1h at 60°C and deparaffinized with a series of xylene and ethanol washes. Once dried for 5min at 60°C, slides were incubated with RNAscope hydrogen peroxide for 10min at room temperature. Target retrieval was then performed for 10min in RNAscope target retrieval solution using a steamer constantly held at 94-95°C. A hydrophobic barrier was drawn on each slide and let dry overnight at room temperature. The next day, protease treatment was performed for 30min at 40°C in a HybEZ Oven (Advanced Cell Diagnostics). Afterward, slides were incubated for 2h at 40°C with the following RNA scope probes: Hs-LETR1-C1 (563721-C1, Advanced Cell Diagnostics), Hs-Prox1-C3 (530241-C3, Advanced Cell Diagnostics; as lymphatic marker), Hs-Malat-1-C2 (400811-C2, Advanced Cell Diagnostics; as positive control, not shown), 3-Plex Negative Control Probe (320871, Advanced Cell Diagnostics; not shown). Slides were then incubated with a series of RNAscope amplifiers, and HRP-channels were developed accordingly to RNA FISH Multiplex Fluorescent v2 Assay user manual. LETR1 and Malat-1 RNAscope probes were visualized by Opal 570 reagents (1:1500 dilution, Perkin Elmer); Prox-1 probes were visualized by Opal 650 reagents (1:1500 dilution, Perkin Elmer). Slides were additionally co-stained with rabbit anti-human vWF polyclonal antibody (Dako) at a dilution of 1:100 overnight at 4°C followed by donkey Alexa488-conjugated anti-rabbit secondary antibody (Thermo Fisher Scientific) at a dilution of 1:200 for 30min at room temperature. All slides were finally counterstained with Hoechst dye diluted 1:10000 in DPBS for 5min at room temperature and mounted with proLong gold antifade mountant (Thermo Fisher Scientific). Z-stacks of fluorescence images spanning over the entire tissue section were acquired using an inverted confocal microscope (Zeiss LSM 880). Z-projection of acquired images was done using ImageJ (ver. 2.0.0-rc-69/1.52i)^97^.

### FACS sorting of primary BECs and LECs followed by qPCR

Single-cell suspensions from skin samples of adult subjects were prepared as described above. Subsequently, isolated single cells were stained with mouse anti-human CD34 biotinylated antibody (clone 581, Thermo Fisher Scientific) diluted 5*µ*L for 10^6^ cells in FACS buffer (DPBS with 2% FBS and 1mM EDTA) for 30min at 4°C. After washing once with FACS buffer, isolated single cells were co-stained in FACS buffer with FITC-conjugated mouse anti-human CD45 antibody (1:25; clone HI30, Biolegend), PE-conjugated mouse anti-human CD31 antibody (1:25; clone WM59, BD Pharmingen), PerCP-conjugated streptavidin (1:400; Biolegend), Zombie NIR (1:500; BioLegend) for 30min at 4°C. After washing in FACS buffer, isolated single cells were filtered and sorted for living, CD45-CD31+ CD34low (LEC), or CD34 high (BEC) directly into test tubes containing 250*µ*L RLT plus lysis buffer, using a FACSAria (BD Biosciences). RNA was isolated using the RNeasy Plus Micro kit (Qiagen), according to the manufacturer’s instructions, including DNase digestion. qPCR was performed on an HT7900 system and analyzed as described above.

### Chromatin isolation by RNA purification followed by sequencing (ChIRP-Seq)

Chromatin isolation by RNA purification followed by DNA sequencing (ChIRP-Seq) was performed as previously described^54^ with minor modifications. Briefly, ChIRP probes (37 x 19-20 nucleotides) targeting LETR1 were designed using the Stellaris Probe Designer (LGC Biosearch Technologies). As non-specific control, 17 probes targeting the bacterial gene LacZ were selected from^54^. Probes were randomly biotinylated using the Photoprobe Labelling Reaction kit (Vector Lab), according to the manufacturer’s instructions. Probe concentrations were determined by Nanodrop. Probes targeting LETR1 were divided into odd and even sets to be used as an additional internal control. Probe sequences are listed in Supplementary Table 8. A total of 60 million LECs per replicate were cross-linked using 1% glutaraldehyde (Sigma) for 10min at room temperature. Crosslinking was quenched with 0.125M glycine (Sigma) for 5min at RT. LECs were rinsed with PBS (Ambion) twice and pelleted for 5min at 2,000rpm. Between 60-80 mg pellets were lysed in 1mL lysis buffer (50mMTris-Cl pH 7.0, 10mM EDTA, 1%SDS, 1mM PMSF, complete protease inhibitor cocktail (Roche), 0.1U/mL RiboLock RNase inhibitor (Thermo Fisher Scientific)) and the cell suspension was sonicated for 1.5-2h until DNA was in the size range of 100-500bp using the Covaris S220 system (Covaris, Peak Power: 140, Duty Factor: 5.0%, Cycles per Burst: 200) with the following on-off intervals: 5×4min, 1×10min, and 4-6×15min. At this point, the sonicated chromatin was divided into 3 equal samples of 1mL (LETR1- Odd, LETR1-Even, and LacZ) and each sample was diluted with 2mL hybridization buffer (750mM NaCl, 1%SDS, 50mM Tris-Cl pH 7.0, 1mM EDTA, 15% formamide, 1mM PMSF, complete protease inhibitor cocktail, 0.1U/mL RiboLock RNase inhibitor). Two aliquots of 20*µ*L chromatin were used as input RNA and DNA. Diluted chromatin samples were then incubated with 100pmol probes and mixed by rotation at 37°C overnight. After overnight hybridization, the three samples (LETR1-Odd, LETR1-Even, and LacZ) were pulled down using 120*µ*L Dynabeads M-270 streptavidin magnetic beads (Thermo Fisher Scientific) for 30min at 37°C with rotation. After five washes with washing buffer (2xSSC, 0.5% SDS, 1mM PMSF), 100*µ*L beads were used for RNA isolation and 900*µ*L for DNA isolation. RNA aliquots were incubated in 80*µ*L RNA proteinase K solution (100mM NaCl, 10mM Tris-Cl pH 7.0, 1mM EDTA, 0.5% SDS, 0.2U/*µ*L proteinase K (Ambion)) for 45min at 50°C with rotation and boiled for 10min at 95°C. 500*µ*L Qiazol (Qiagen) were added to each sample, and RNA was extracted according to the manufacturer’s instructions of the miRNeasy mini kit (Qiagen). Equal amounts of isolated RNA (10ng) were subjected to one-step real-time PCR using the One-Step SYBR PrimeScript RT-PCR kit (Takeda Bio) on an HT7900 system in order to determine the percentage of RNA retrieval. DNA, on the other hand, was eluted twice using 150*µ*L DNA elution buffer (50mM NaHCO3, 1% SDS, 200mM NaCl, 1mM PMSF, 100*µ*g/mL RNase A (Thermo Fisher Scientific), 100U/mL RNase H (New England BioLabs)) and incubated for 30min at 37°C with shaking. DNA eluates were combined and incubated with 300*µ*g proteinase K for 45min at 50°C. DNA was purified using phenol:chloroform:isoamyl (25:24:1, Thermo Fisher Scientific) followed by the MiniElute PCR purification kit (Qiagen). The quality of DNA was assessed using a 2100 Bioanalyzer instrument (Agilent). Finally, DNA was used to prepare libraries using the ThruPLEX DNA-Seq kit (Rubicon), and sequencing was performed according to the manufacturer’s protocol on a HiSeq system (Illumina).

### ChIRP-Seq analysis

Analysis including alignment and peak calling was performed following the published ChIRP-Seq pipeline from the Chang Lab at Stanford University^54^ with minor modifications. Briefly, raw reads were firstly trimmed using SolexaQA (ver. 3.1.7.1)^100^ with -dynamictrim and -lengthsort. Trimmed reads were then uniquely mapped to human reference genome hg38 using bowtie (ver. 1.1.1)^101^. Mapping parameters were -m 1 -chunkmbs 1024 -p 6. Peaks against input were called using MACS 2.0 (ver. 2.1.1)^102^ with following settings callpeak -f SAM -B --SPMR -g hs --bw 200 -m 10 50 -q 0.01. Finally, peaks were filtered based on fold enrichment against input lane > 3, Pearson correlation > 0.2, and average coverage > 1.25. ChIRP peaks are listed in Supplementary Table 9. Genomic location analysis of significant peaks was performed using bedtools (ver. 2.27.1)^103^ and FANTOM CAT annotations^13^ for gene body, exon, and promoter. Enrichment analyses of protein-coding genes, as well as of common genes between LETR1-ASOKD and ChIRP-Seq, were performed using SuperExactTest (ver. 1.0.0)^91^. As background, total genes annotated in the FANTOM CAT database^13^ (n = 124,047) and total genes from ChIRP and ASOKD data (n = 13,127) were respectively used. The circular plot of the identified gene targets was generated using Circos (ver. 0.69-7)^104^.

### Cloning of LETR1 and SEMA3C, and lentivirus production

For cloning, full-length LETR1 transcript (LETR1-1; most abundant RACE transcript) and the coding sequences (CDSs) of SEMA3C (ENST00000265361.7) were PCR amplified from neonatal LEC cDNA using Phusion high-fidelity DNA polymerase (New England BioLabs), according to the manufacturer’s protocol. Amplified fragments were digested with BamHI and NotI restriction enzymes (New England BioLabs) overnight at 37°C and cloned into a linearized pCDH-EF1alpha-MCS-BGH-PGK-GFP-T2A-Puro lentivector (CD550A-1, System Biosciences) using T4 DNA ligase (New England BioLabs) for 20min at room temperature. Plasmids were transformed, isolated, and checked, as described above. The empty pCDH plasmid was used as a negative control. PCR cloning primers are listed in Supplementary Table 11. Lentivirus production for each vector was carried out as described above.

### Establishment of LECs overexpressing LETR1 and SEMA3C

1.2×10^5^ LECs per well were seeded into a 6-well plate and cultured overnight. LECs were then infected with medium containing viruses overexpressing LETR1 and SEMA3C diluted at a 10 MOI and 10*µ*g/mL polybrene (hexadimethrine bromide). Plates were then sealed with parafilm and centrifuged at 1,250rpm for 1.5h at room temperature. After 16-24h, the virus-containing medium was changed. 24h later, infected LECs were subjected to antibiotic selection using puromycin at a 1*µ*g/mL concentration (Thermo Fisher Scientific). Once plates were confluent, infected LECs were split into 10cm dishes at 3×10^5^ cells per dish and further selected with 2-5*µ*g/mL puromycin until full confluence was reached. Finally, infected LECs were aliquoted and frozen down for further experiments. To check RNA levels, approx. 70’000 infected LECs were lysed, total RNA was extracted, and cDNA synthesis and qPCR were performed as described above. To check protein levels, a confluent 6cm dish of infected LECs was lysed in 350*µ*L lysis buffer (25mM HEPES, 5mM EDTA, 1% triton-X, 150mM NaCl, 10% glycerol, complete protease inhibitor cocktail). The lysate was then centrifuged at 13,000rpm for 20min at 4°C, and the supernatant was collected in a 1.5mL Eppendorf tube. SEMA3C protein levels were checked by western blot as described above using rabbit anti-human SEMA3C antibody (1:1000, Thermo Fisher Scientific). Beta-actin (housekeeping gene) was detected with a rabbit anti-beta-actin antibody (1:5000, Abcam) (Supplementary Figure 6b).

### Rescue of cell growth phenotype experiments

For LETR1, 5×10^5^ LECs infected with pCDH-EV or pCDH-LETR1 were seeded into 10cm dishes and cultured overnight in starvation medium (EBM supplemented with 1% FBS). The next day, infected LECs were transfected with scramble ASO or LETR1-ASO2 for 24h as described above.

For KLF4, a consecutive transfection of ASOs and siRNAs was performed. siRNA sequences are listed in Supplementary Table 11. 350’000 LECs were seeded into 10cm dishes and cultured overnight in starvation medium. The next day, LECs were transfected with scramble control ASO or LETR1-ASO2, as described above. 24h post-transfection with ASOs, the medium was changed, and LECs were treated for additional 24h with 20nM of high GC scramble control siRNA or two siRNAs targeting KLF4 (Thermo Fisher Scientific) and 32*µ*L Lipofectamine RNAiMAX previously mixed in 800*µ*L Opti-MEM.

At the end of the experiment, LECs were detached and collected. 1.5×10^5^ LECs per replicate were then transferred in a 96 U-bottom plate (Greiner bio-one). Aliquots of approx. 1×10^5^ LECs were lysed and subjected to qPCR, as described above. Cell cycle analysis using flow cytometry was performed as described above. To detect Ki-67, an eFluor 450-conjugated rat anti-human Ki-67 antibody (clone: SolA15, ebioscience) was used at a 1:200 dilution.

### Rescue of cell migration phenotype experiments

2.5×10^5^ LECs infected with pCDH-empty vector (EV), pCDH-LETR1, and pCDH-SEMA3C were seeded into 6cm dishes and cultured for at least 6h. Infected LECs were then transfected with 20nM of scramble control ASO or LETR1-ASO2 and 8*µ*L Lipofectamine RNAiMAX previously mixed in 800*µ*L Opti-MEM according to the manufacturer’s instructions. 24h post-transfection, LECs were detached, and 15,000 were seeded into a 96-well plate and cultured overnight. The next day, infected LECs were pre-treated with proliferation inhibitor, scratched, and monitored for cell migration as described above.

### Biotin-RNA pull-down followed by mass spectrometry

Biotin-RNA pull-down experiments were performed as previously described^57^ with minor modifications. To prepare nuclear lysate, 40 million LECs per replicate were collected and washed once with DPBS. The cell pellet was resuspended in 40mL consisting of 8mL DPBS, 24mL RNase-free H_2_O, and 8mL nuclear isolation buffer (1.28M sucrose, 40mM Tris-HCl pH 7.5, 20mM MgCl_2_, 4% triton X-100). After mixing by inversion, LECs were incubated on ice for 20min with occasional mixing. Cells were then centrifuged at 1,668rpm for 15min at 4°C. The nuclear pellet was resuspended in 2mL of RNA pull-down buffer (150mM KCl, 2.5mM MgCl_2_, 25mM Tris-HCl pH 7.5, 5mM EDTA pH 8, 0.5% NP-40, 0.5mM DTT, complete protease inhibitor cocktail, 100U/mL Ribolock RNase inhibitor). Lysed nuclei were transferred into a 7mL Dounce homogenizer (Kimble) and sheared mechanically using 30-40 strokes with pestle B. Next, the nuclear lysate was transferred to a fresh tube and sonicated 2x 30s at a high intensity (50% cycle and 90% power) using a Sonopuls HD2070 (Bandelin). 10U/mL DNase I (Thermo Fisher Scientific) were subsequently added to the nuclear lysate and incubated for 30min at 4°C while rotating. The nuclear lysate was further sonicated for another 2x 30s at high intensity. The nuclear lysate was centrifuged at 13,000rpm for 10min at 4°C. Finally, the supernatant was collected into a fresh tube, and glycerol was added to reach a 10% final concentration. Resulting clear nuclear lysate was flash-frozen in liquid nitrogen and stored at −80°C for later use. Nuclear fractionation was checked after performing western blot analysis, as described above, of GAPDH (cytoplasmic protein) and Histone H3 (nuclear protein) using rabbit anti-GAPDH antibody (1:5000, Sigma) and rabbit anti-histone H3 antibody (1:10000, Sigma) antibodies (Supplementary Figure 7a).

To produce biotinylated RNA, full-length LETR1-1, determined by RACE (Supplementary Table 7), and antisense strand were cloned into pcDNA3.1 backbone. Both transcripts were amplified by PCR using Phusion high-fidelity DNA polymerase (New England BioLabs), according to the manufacturer’s protocol. Amplified fragments were digested with BamHI and XbaI restriction enzymes (New England BioLabs) overnight at 37°C. After gel purification, digested fragments were cloned into a linearized pcDNA3.1 backbone (from 64599, Addgene) using T4 DNA ligase for 20min at room temperature. Cloned vectors were transformed into one shot TOP10 chemically competent cells, according to the manufacturer’s instructions. Plasmids were isolated using the Nucleospin Plasmid kit (Machery Nagel). Sequences of inserted fragments were checked by Sanger sequencing (Microsynth). PCR cloning primers are listed in Supplementary Table 11. Subsequently, both transcripts were biotin-labeled after in vitro transcription from 1*µ*g linearized pcDNA3.1-LETR1-1 and pcDNA3.1-LETR1-1-antisense plasmids for 1h at 37°C using Ampliscribe T7-flash biotin-RNA kit (Lucigen). Biotinylated LETR1 sense and antisense RNA were then treated with RNase-free DNase I for additional 15min at 37°C. Both biotinylated RNAs were purified by ammonium acetate precipitation, as described by the manufacturer. After determining the concentration using Nanodrop 1000, the integrities of sense and antisense LETR1 transcripts were tested by gel electrophoresis (Supplementary Figure 7b).

To perform RNA pull-down, 150*µ*L Dynabeads M-270 streptavidin magnetic beads were washed twice with RNA pull-down buffer. For each condition, 60*µ*L washed beads were then incubated with 1.5mg nuclear lysate for 30min at 4°C. During nuclear pre-clearing, 100pmol per condition of biotinylated RNAs were denatured by heating to 65°C for 10min and cooled down slowly to 4°C. Pre-cleared nuclear extract was further diluted to 2mL using RNA pull-down buffer and incubated with 100pmol biotinylated RNA for 1h at 4°C on a rotatory shaker. Next, 60*µ*L washed streptavidin magnetic beads were added and further incubated for 45min at 4°C. Beads were carefully washed five times in RNA pull-down buffer. Bound proteins were finally eluted twice by adding 3mM biotin in PBS (Ambion) to the beads and incubating them for 20min at room temperature and for 10min at 65°C. Eluted proteins were subjected to protein identification by mass spectrometry at the Functional Genomics Center Zurich (FGCZ). Proteins were pelleted by TCA precipitation using 20% TCA. Protein pellets were washed twice with cold acetone. Dry pellets were then dissolved in 45*µ*L trypsin buffer (10mM Tris, 2mM CaCl2, pH 8.2) plus 5*µ*L trypsin (100ng/*µ*L in 10mM HCl) and 1.5*µ*L 1M Tris pH 8.2. After microwaving for 30min at 60°C, dried samples were dissolved in 0.1% formic acid. Digested peptides were finally analyzed by LC/MS.

### Analysis of RNA-pull down data

Database searches were performed using Mascot software (Matrix Science). Search results were then loaded into Scaffold software (ver. 4.8.7) to perform statistical analysis. Only proteins with 1% protein FDR, a minimum of 2 peptides per protein, 0.1% peptide FDR, and present in both LETR1 replicates but not in the antisense control were considered. PPI network for the proteins identified by RNA-biotin pull-down was generated using the STRING web tool (https://string-db.org/cgi/input.pl)^58^. The human PPI database was used for the analysis, while default values were used for the rest of the parameters. The identified proteins are listed in Supplementary Table 10.

### Native RNA immunoprecipitation followed by qPCR

RNA immunoprecipitation experiments were performed as previously described^105^ with minor modifications. To prepare nuclear lysate, 40 million LECs per replicate were collected and washed once with DPBS. The cell pellet was resuspended in 40mL consisting of 8mL DPBS, 24mL RNase-free H_2_O, and 8mL nuclear isolation buffer (1.28M Sucrose, 40mM Tris-HCl pH 7.5, 20mM MgCl_2_, 4% triton X-100). After mixing by inversion, LECs were let stand on ice for 20min with occasional mixing. Cells were then centrifuged at 1,668rpm for 15min at 4°C. Nuclear pellet was resuspended in 2mL of RIP buffer (150mM KCl, 25mM Tris-HCl pH 7.5, 5mM EDTA pH 8, 0.5% NP-40, 0.5mM DTT, complete protease inhibitor cocktail, 100U/mL Ribolock RNase inhibitor) and incubated for 5min on ice. Lysed nuclei were transferred into a 7mL Dounce homogenizer (Kimble) and sheared mechanically using 30-40 strokes with pestle B. Next, the nuclear lysate was transferred to a fresh tube and centrifuged at 13,000rpm for 10min at 4°C. Finally, the supernatant was collected into a fresh tube, and glycerol was added to reach a 10% final concentration. The resulting clear nuclear lysate was snap-frozen in liquid nitrogen and stored at −80°C for later use. For each replicate, 1mg nuclear lysate in 1.5mL RIP Buffer was incubated with 2.5*µ*g rabbit anti-human RBBP7 antibody (Cell Signaling) or rabbit IgG isotype control antibody (I5006, Sigma) overnight at 4°C with gentle rotation. Two aliquots of 15*µ*L nuclear lysate were used as input RNA and protein. Next, 50*µ*L washed Dynabeads protein A magnetic beads (Thermo Fisher Scientific) were added to each sample and incubated for 2h at 4°C. After five washes with 500*µ*L RIP buffer, each sample’s beads were resuspended in 500*µ*L RIP buffer. 25*µ*L beads of each sample were then transferred to a fresh tube and subjected to western blot analysis as described above using the same RBBP7 antibody at a dilution of 1:1000 (Figure 7c). In order to prevent masking from denatured IgG heavy chains, we detected the primary antibody with a mouse anti-rabbit IgG conformation-specific secondary antibody (Cell Signaling). The rest of the sample’s beads were resuspended in 700*µ*L Qiazol (Qiagen), and RNA was extracted using the miRNeasy mini kit, according to the manufacturer’s instructions. Isolated RNA was then subjected to cDNA synthesis and qPCR, as described above. LETR1 Ct values were normalized to the housekeeping gene GAPDH.

### Statistical analysis

All statistical analyses were performed using GraphPad Prism software (ver. 7.0.0). P-values were calculated after performing ordinary and RM two- or one-way ANOVA with Dunnett’s correction, paired or unpaired Student’s *t*-test as indicated. Statistical significance was determined when P < 0.05. If not alternatively specified, all error bars represent mean values with SD.

## Data availability statement

Further information and requests for resources and reagents should be directed to and will be fulfilled by Michael Detmar. All software used in this study are specified in the related method paragraph with the corresponding version. All unprocessed sequencing data are going to be deposited in DDBJ DRA (https://www.ddbj.nig.ac.jp/dra/index-e.html) and on the FANTOM consortium website (https://fantom.gsc.riken.jp/6/). LC/MS data are available at the Functional Genomic Center Zurich (Yolanda Auchli; ID:11594).

## Acknowledgments

This study was financially supported by the ETH Zurich (grant ETH-24 171), the Swiss National Science Foundation (grant 310030_166490), and the European Research Council (advance grant LYVICAM). We acknowledge Jeannette Scholl for her support in performing the smRNAFISH expression analysis, as well as Dr. Raffaella Santoro and Dominik Bär, University of Zurich, for crucial advice on the RNA pull-down assay. Finally, we also thank all the members of the FANTOM6 project for fruitful discussions and support throughout the project.

## Author Contributions

L.D. design the project, performed the *in silico* analyses and wet lab experiments, and wrote the manuscript. S.A. performed *in silico* analysis of ASO-mediated knockdown CAGE sequencing data, helped analyze the RNA immunoprecipitation data, discussed and interpreted the results, and strongly contributed to figures, general discussion and writing of the manuscript. E.S. helped performing *in vitro* experiments, established the cell cycle progression method, and strongly contributed to general discussions and writing of the manuscript. C.T. isolated the adult LECs and BECs, performed the FACS sorting of *ex vivo* LECs and BECs, and strongly contributed to experimental design, general discussion and comments on the manuscript. T.K. helped with the ChIRP-Seq experiment by performing the library preparation and sequencing, contributed to general discussions and comments for the manuscript. C-C.H. helped analyze the RNA-Seq and CAGE-Seq data of LECs and BECs, and contributed to general discussions and comments for the manuscript. S.D.B. helped performing *in vitro* studies and contributed to general discussions as well as added comments to the manuscript. D.M. performed the subcellular fractionation of lncRNA targets in LECs and BECs and provided comments on the manuscript. Y.H. supported L.D. in the ChIRP-Seq analysis, contributed to general bioinformatics discussions, helped interpreting the results, and provided comments for the manuscript. M.Da. performed lineage check of LECs and BECs and isolated RNA to be subjected to both RNA-Seq and CAGE-Seq, and provided comments for the manuscript. L.Di. made crucial contributions to *in vitro* as well as general experimental design, helped interpret the results, and contributed in writing the manuscript. P.C. and M.J.L.dH. discussed and interpreted the results, provided crucial comments for the progress of the project, and revised the manuscript. J.W.S. and M.D. supervised and guided the entire project, provided resources for all the experiments, helped in interpreting the results and writing of the manuscript.

