## Supplemental Figures 1-8 for "LETR1 is a lymphatic endothelial-specific lncRNA that governs cell proliferation and migration through KLF4 and SEMA3C"

Supplementary Figure 1

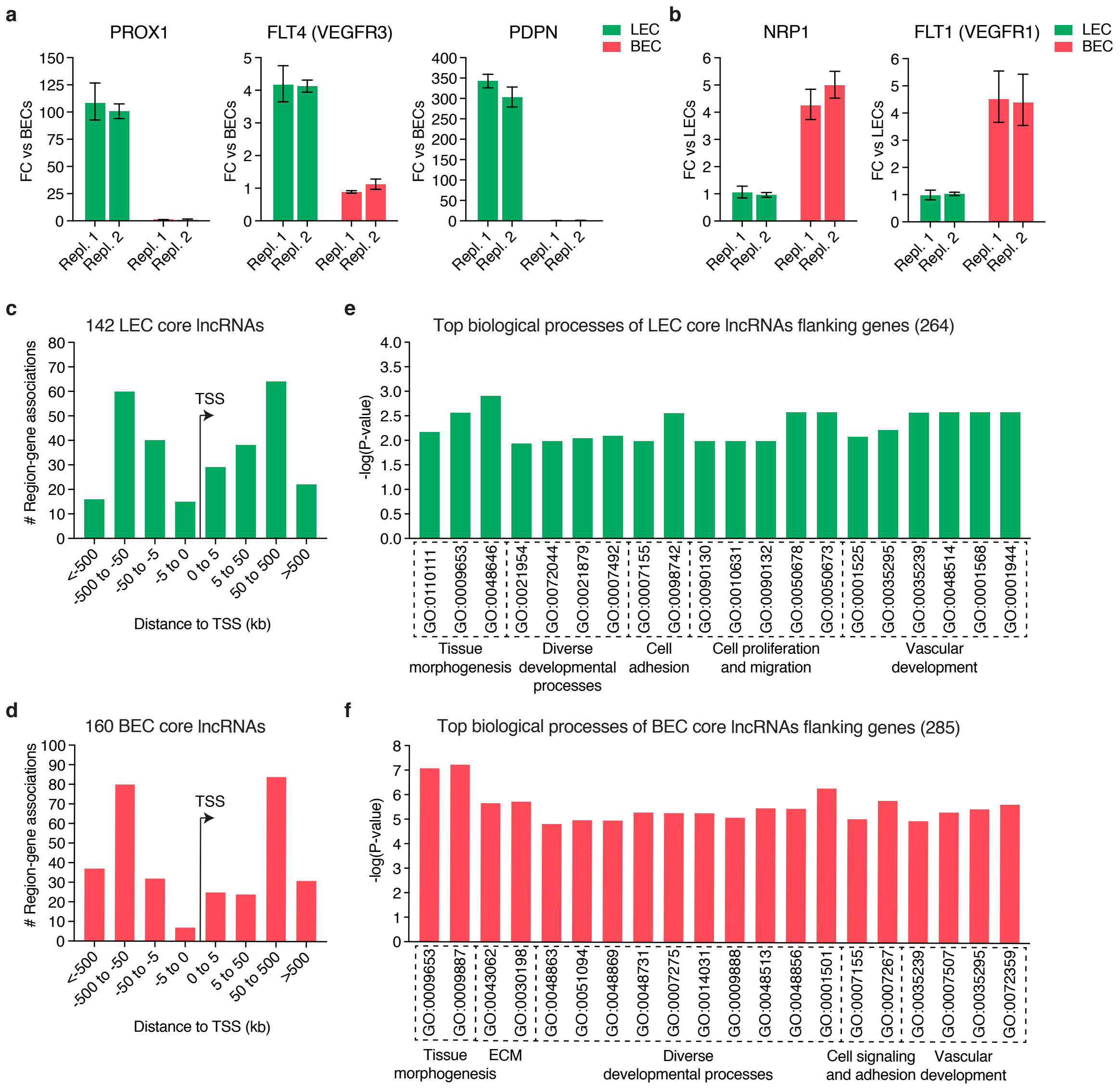

**Supplementary Figure 1: Confirmation of LEC and BEC identity and analysis of BEC and LEC core lncRNA flanking protein-coding genes.**

**(a, b)** Validation of LEC and BEC identities through analysis of LEC (a, PROX-1, FLT4, PDPN) and BEC (b, NRP1, FLT1) specific markers using qPCR. Bars represent fold change (FC) to either average LEC or BEC expression as mean + upper and lower error (n = 3 technical replicates). RPLP0 was used as the housekeeping gene.

**(c, d)** Distribution of flanking genes determined with GREAT[1] of LEC (c) and BEC (d) core lncRNAs by the use of the association rule "two nearest genes" with a maximal extension from the lncRNA transcriptional start sites (TSS) of 10 Mb.

**(e, f)** Top significant (P-value < 0.05) enriched biological processes of LEC (e) and BEC (f) core lncRNA flanking genes, using gProfileR package[2] (relative depth 2-5). Terms were manually ordered according to their biological meaning. Only genes with expression values (TPM & CPM) > 0.5 in LECs or BECs were used as background.

Supplementary Figure 2

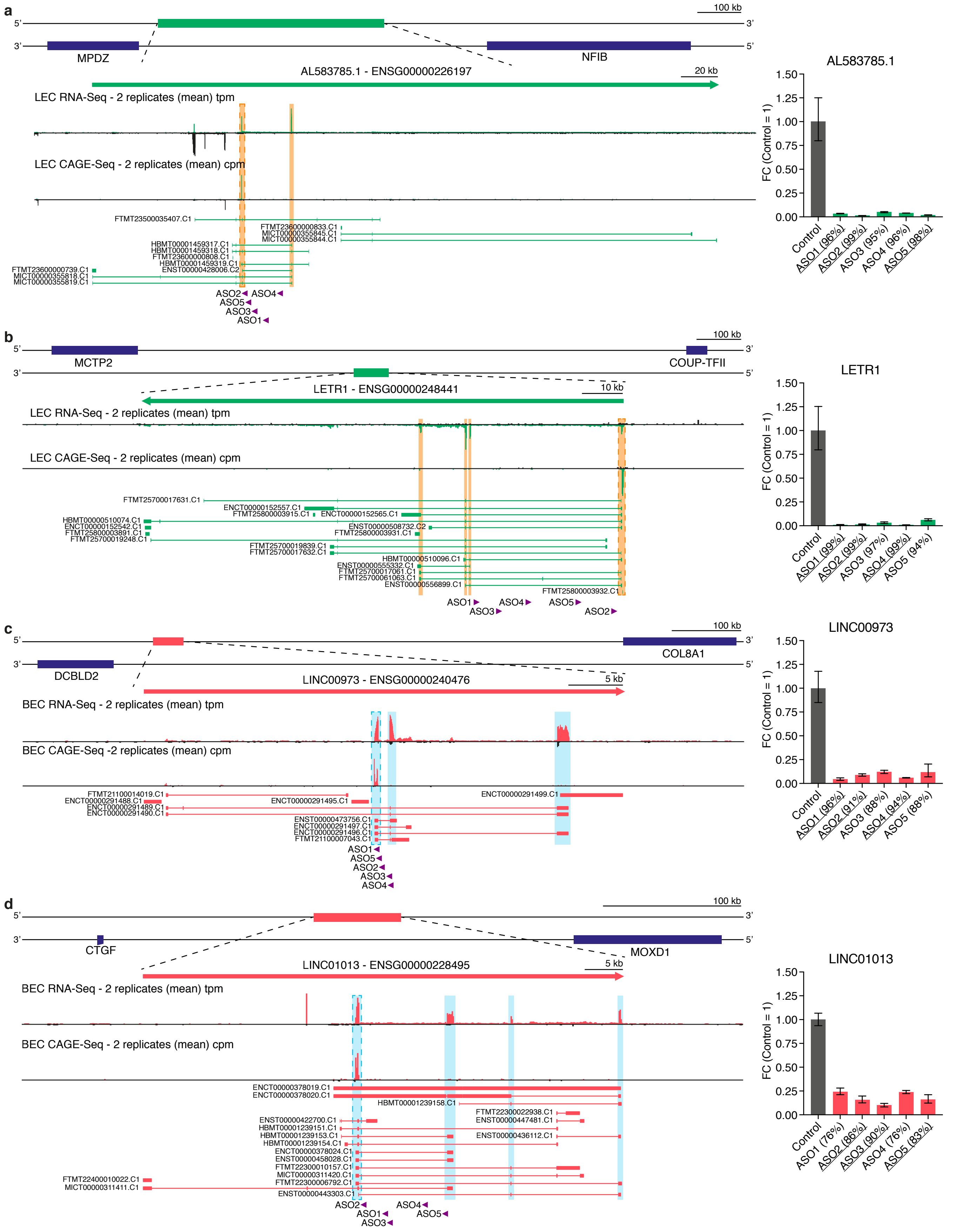

**Supplementary Figure 2: ASO design strategy for LEC and BEC lncRNA candidates and selection of 3 most efficient ASOs per target through qPCR.**

**(a-d)** Schematic representation of the genomic regions of 2 LEC (a, b) and 2 BEC (c, d) lncRNA candidates with their flanking genes (blue boxes) according to FANTOM CAT database[3]. Magnifications show lncRNA gene region with respective RNA-Seq (TPM, 2 replicates) and CAGE-Seq (CPM, 2 replicates) signals in LECs (in green) or BECs (in red), related transcripts (green: LEC; red: BEC), and ASO locations (in purple). RNA-Seq and CAGE-Seq signals were visualized through the Zenbu genome browser[4]. Orange/cyan-dashed boxes represent the overlap between RNA-Seq and CAGE-Seq peaks. Bar charts show knockdown efficiencies determined by qPCR after 48h transfection of five ASOs targeting the 2 LEC (a, b) and the 2 BEC (c, d) lncRNAs in neonatal LECs or BECs derived from the same donor. Selected ASOs for CAGE-Seq are underlined. Bars represent FC compared to control ASO (FC = 1) as mean ± upper and lower error (n = 3 technical replicates). RPLP0 was used as the housekeeping gene. Percentages of knockdown efficiencies for each ASO are shown in brackets. ASO sequences are listed in Supplementary Table 2.

Supplementary Figure 3

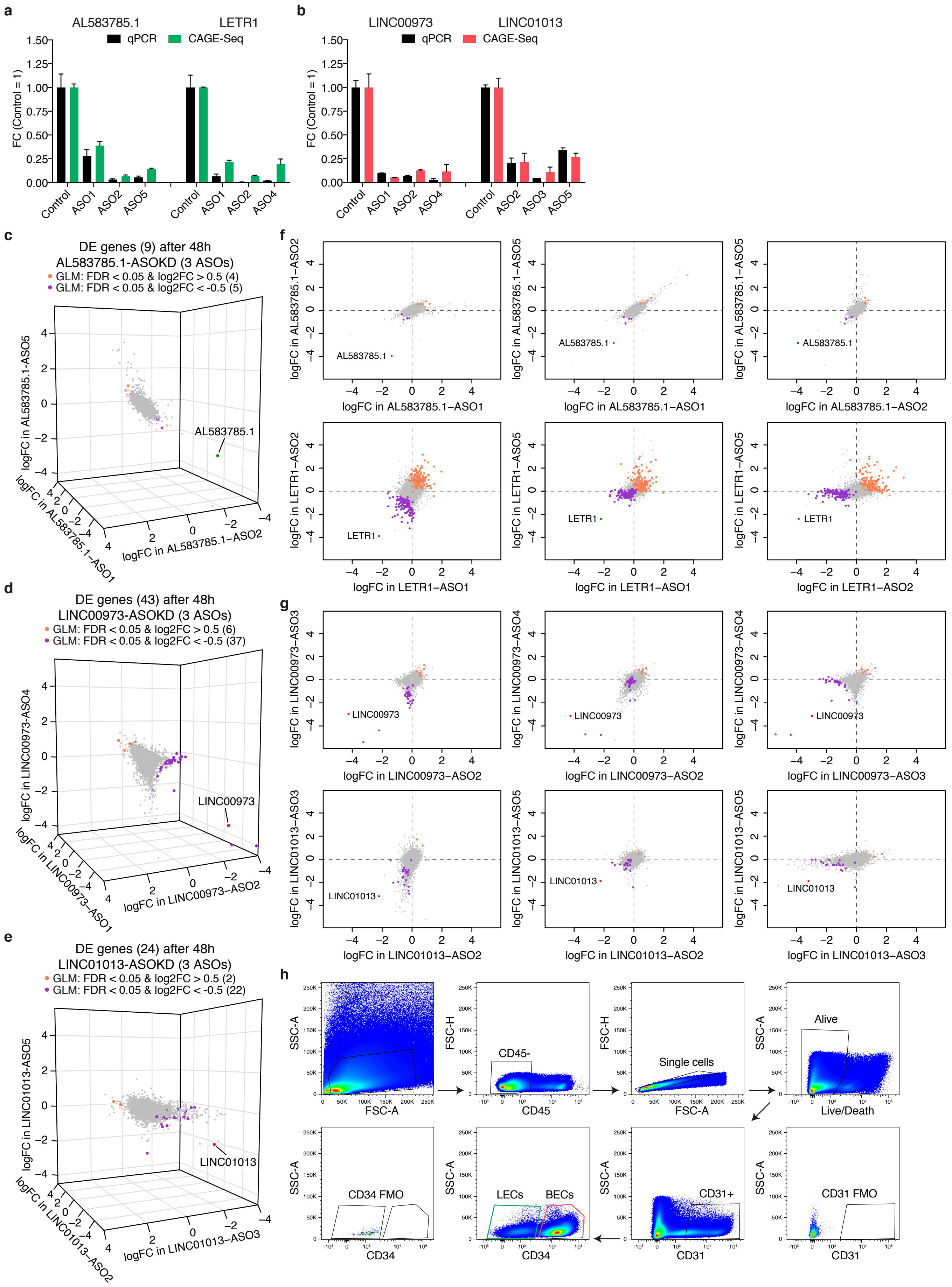

**Supplementary Figure 3: Knockdown efficiency of CAGE-Seq samples, DE analysis after ASOKD of AL583785.1, LETR1, LINC00973, and LINC01013, and gating strategy used to sort LECs and BECs from human skin.**

**(a, b)** Graph showing FC reduction compared to control ASO of LEC (a) and BEC (b) lncRNAs after LETR1-ASOKD determined with qPCR (black bars) and CAGE-Seq (green or red bars). Bars represent mean + SD (n = 2).

**(c-e)** 3-dimensional scatter plots showing log2FC values calculated between single ASO and control ASO through EdgeR[5] for LEC candidate AL583785.1 (c) and BEC candidates LINC00973 (d) and LINC01013 (e). Orange and purple dots represent significantly (FDR < 0.05) up- and downregulated genes (|logFC| > 0.5) after differential expression analysis applying a generalized linear model design (GLM)[5]. Green dot shows AL583785.1; red dots display LINC00973 and LINC01013. DE genes for all lncRNA candidates are listed in Supplementary Table 3.

**(f, g)** 2-dimensional scatter plots showing log2FC values of ASO pairs for each of the 2 LEC (f) and 2 BEC (g) lncRNAs. Orange and purple dots represent up- and downregulated genes. Green dots show AL583785.1 and LETR1; red dots display LINC00973 and LINC01013.

**(h)** Representative flow cytometry plots showing the gating strategy used to isolate LECs and BECs from 3 donors of healthy human skin samples. After gating for living and CD31+ cells, LECs and BECs were sorted based on their CD34 expression. Total RNA was then extracted and subjected to qPCR.

### Supplementary Figure 4

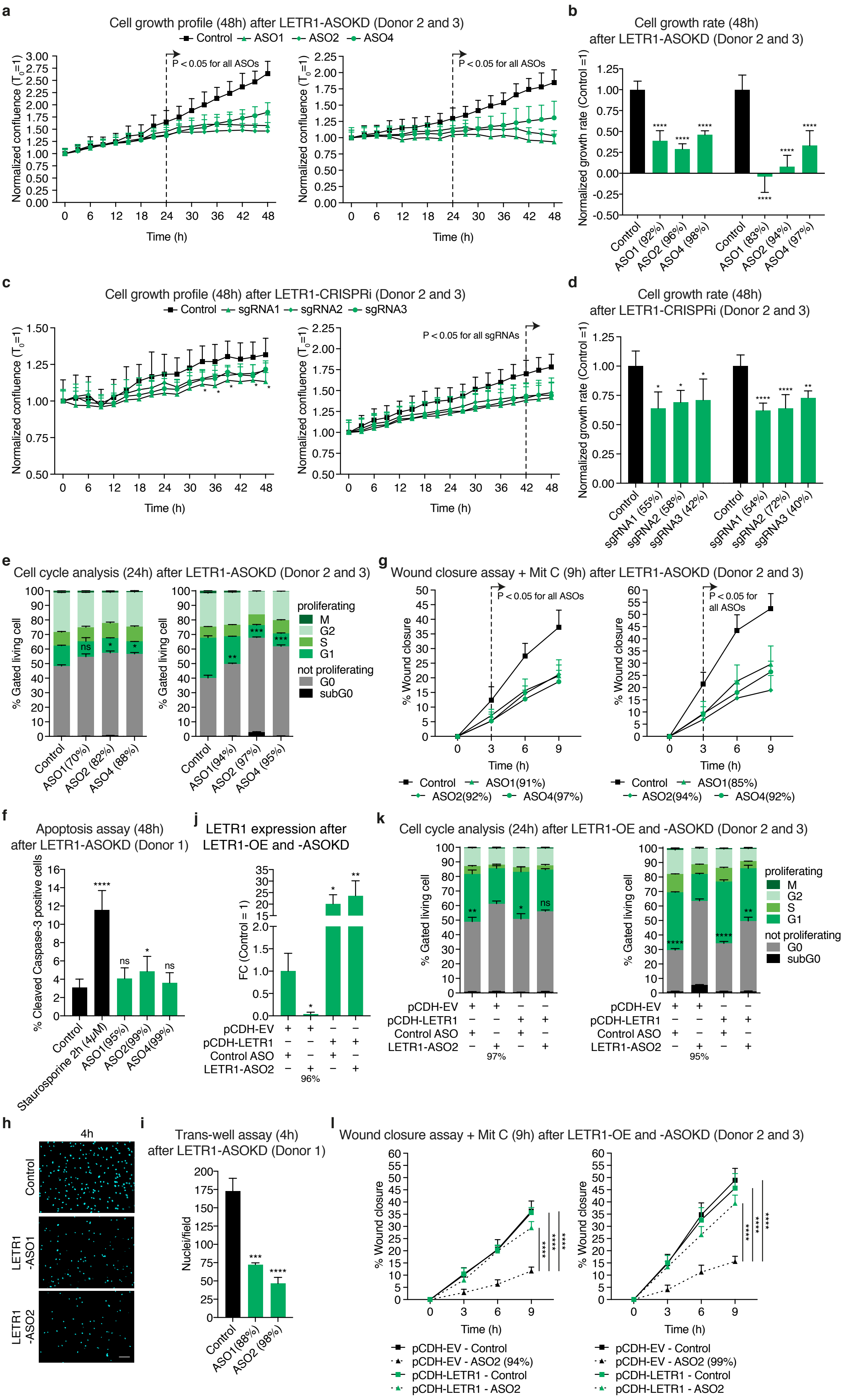

**Supplementary Figure 4: Validation of *in vitro* assays in 2 additional donors of LECs, and results of apoptosis and trans-well assay.**

**(a-d)** Cell growth profiles and normalized growth rates of neonatal LECs derived from 2 additional donors over 48h after ASOKD (a, b) or CRIPSRi-KD (c, d) of LETR1.

**(e)** Quantification of the cell cycle progression analysis of neonatal LECs derived from 2 additional donors after 24h LETR1-ASOKD.

**(f)** Quantification of the apoptosis assay (48h) of neonatal LECs derived from one donor after LETR1-ASOKD. Percentages of cleaved caspase 3-positive cells were determined using ImageJ[6]. Cells incubated for 2h with Staurosporine (4$\mu$M) were used as a positive control.

**(g)** Quantification of the wound closure assay (up to 9h) of neonatal LECs derived from 2 additional donors after LETR1-ASOKD.

**(h)** Representative images of the trans-well assay (4h) of neonatal LECs derived from one donor after LETR1-ASOKD. Nuclei were stained with DAPI. Scale bar represents 120$\mu$m.

**(i)** Quantification of the trans-well assay (4h) of neonatal LECs derived from one donor after LETR1-ASOKD. DAPI-stained nuclei per field were determined using ImageJ[6].

**(j)** qPCR expression levels of LETR1 in pCDH-empty vector (pCDH-EV) and pCDH-LETR1 infected neonatal LECs derived from 3 donors after 24h LETR1-ASOKD. Bars represent FC values against control ASO. GAPDH was used as the housekeeping genes.

**(k)** Quantification of the cell cycle progression analysis of pCDH-EV and pCDH-LETR1 infected neonatal LECs derived from 2 additional donors after 24h LETR1-ASOKD.

**(l)** Quantification of the wound closure assay (up to 9h) of pCDH-EV and pCDH-LETR1 infected neonatal LECs derived from 2 additional donors after LETR1-ASOKD.

Data are displayed as mean + SD (n = 10 in a, b, f, g, and l; n = 5 in c and d; n = 2 in e; n = 3 in i, j, and k). Percentages represent LETR1 knockdown efficiencies after the experiments. *P < 0.05, **P < 0.01, ***P < 0.001, ****P < 0.0001, ns not significant using one-way (for b, d, e, f, i, and k), RM one-way (for j), and two-way (for a, c, g, and l) ANOVA with Dunnett's multiple comparisons test against control ASO/sgRNA, pCDH-EV – Control ASO, or LETR1-ASO2 – control siRNA.

Supplementary Figure 5

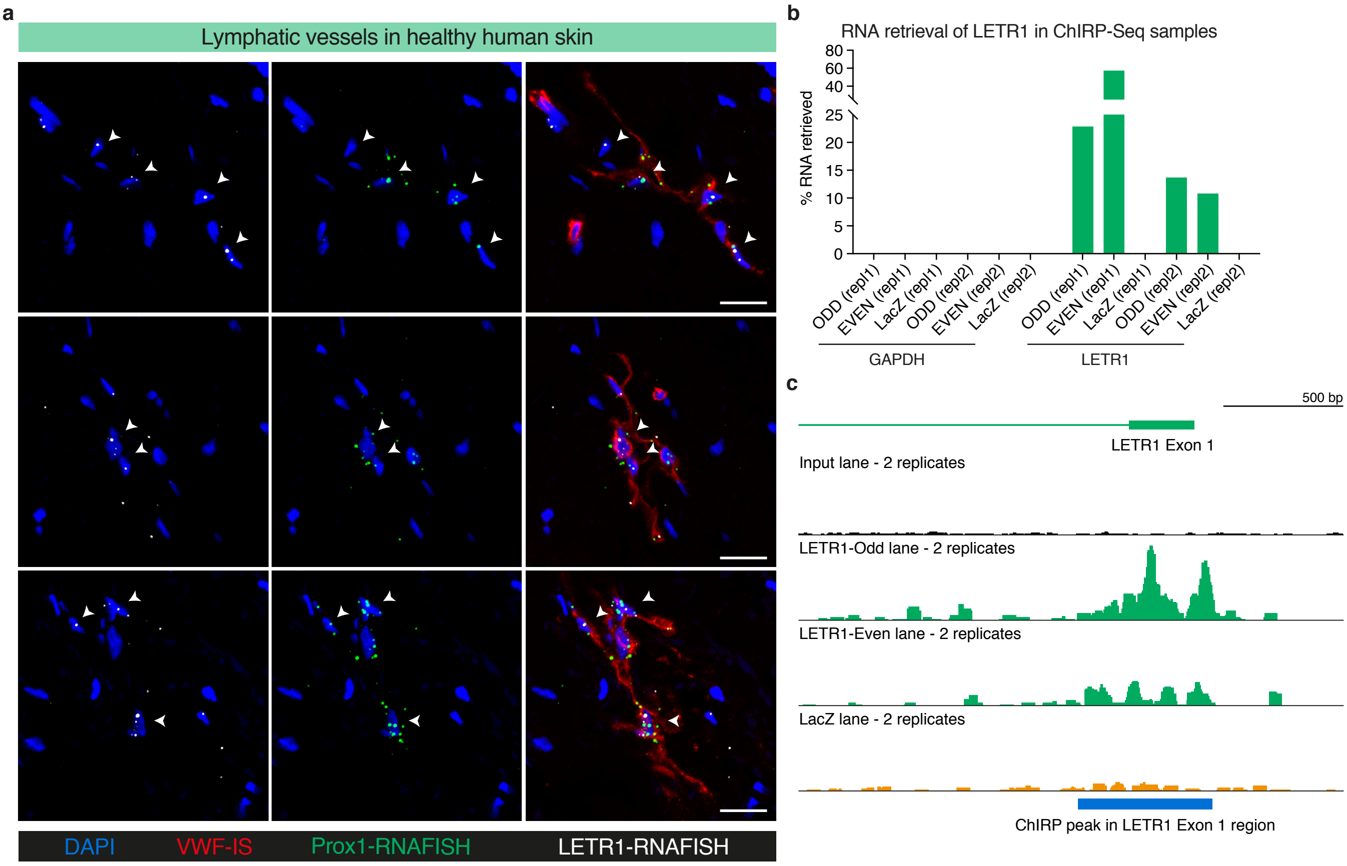

**Supplementary Figure 5: LETR1 *in vivo* expression of LETR1 and results after LETR1 ChIRP-Seq.**

**(a)** Representative images of lymphatic vessels in healthy human skin. Lymphatic vessels were defined as von Willebrand factor (vWF, red – immunostaining) and PROX1 (green – smRNA-FISH) double positive. Scale bars represent 20$\mu$m.

**(b)** RNA retrieval displayed as percentages of GAPDH (negative control) and LETR1 in ODD, EVEN, and LacZ samples (2 replicates of neonatal LECs derived from the same donor). Probe sequences are listed in Supplementary Table 8.

**(c)** Schematic representation of the genomic region of LETR1 – Exon 1 and the corresponding ChIRP-Seq signal of input (black), LETR1-Odd and -Even (green), and LacZ lanes (orange). Significant peak region is shown in blue. ChIRP-Seq signals were visualized through Integrative Genomics Viewer (IGV)[7].

Supplementary Figure 6

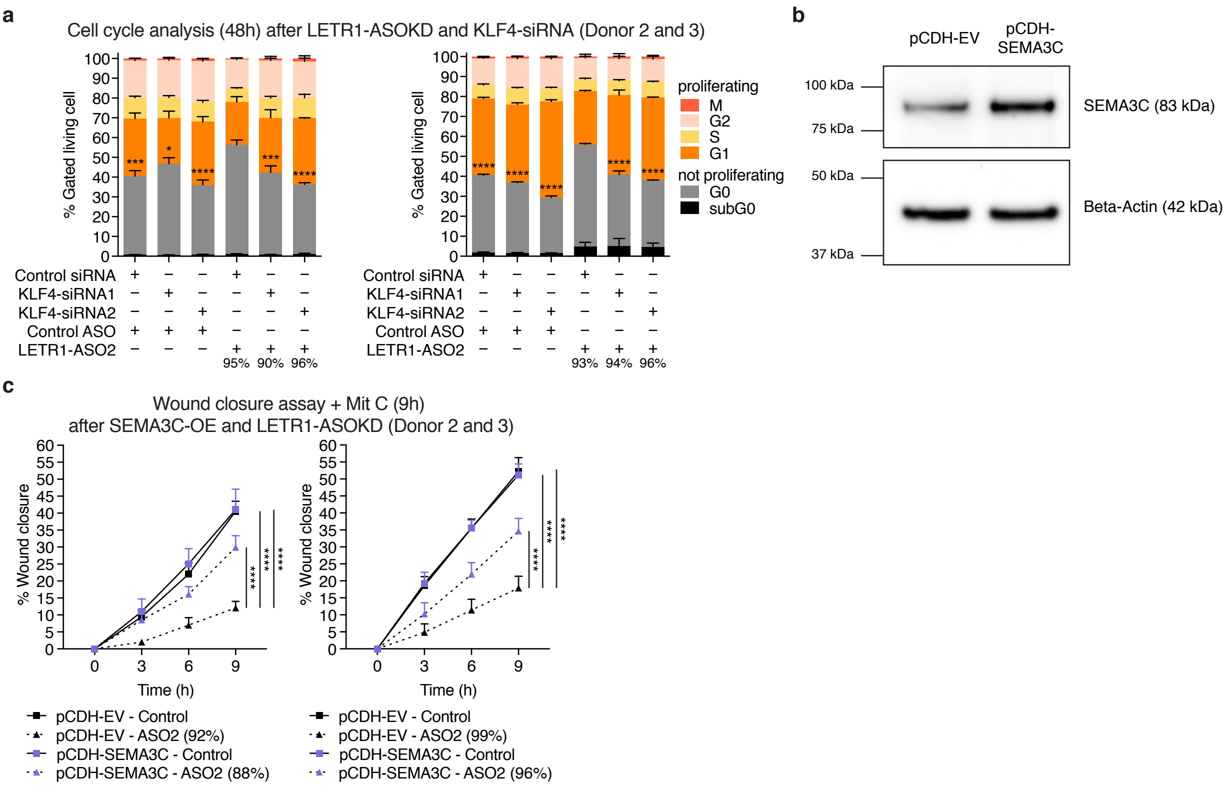

**Supplementary Figure 6: Validation of KLF4 and SEMA3C rescues in 2 additional donors of LECs.**

**(a)** Quantification of the cell cycle progression analysis of neonatal LECs derived from 2 additional donors after 24h LETR1-ASOKD followed by 24h siRNA-KD of KLF4.

**(b)** Western blot images for SEMA3C in pCDH-EV and pCDH-SEMA3C. Uncropped western blot image is shown in Supplementary Figure 8.

**(c)** Quantification of the wound closure assay (up to 9h) of pCDH-EV and pCDH-SEMA3C infected neonatal LECs derived from 2 additional donors after LETR1-ASOKD.

Data are displayed as mean + SD (n = 3 in a; n = 10 in c). Percentages represent the knockdown efficiencies of LETR1 after the experiments. *P < 0.05, ***P < 0.001, ****P < 0.0001 using, ordinary one-way (for a), and two-way (for c) ANOVA with Dunnett's multiple comparisons test against LETR1-ASO2 – control siRNA or pCDH-EV – ASO2.

Supplementary Figure 7

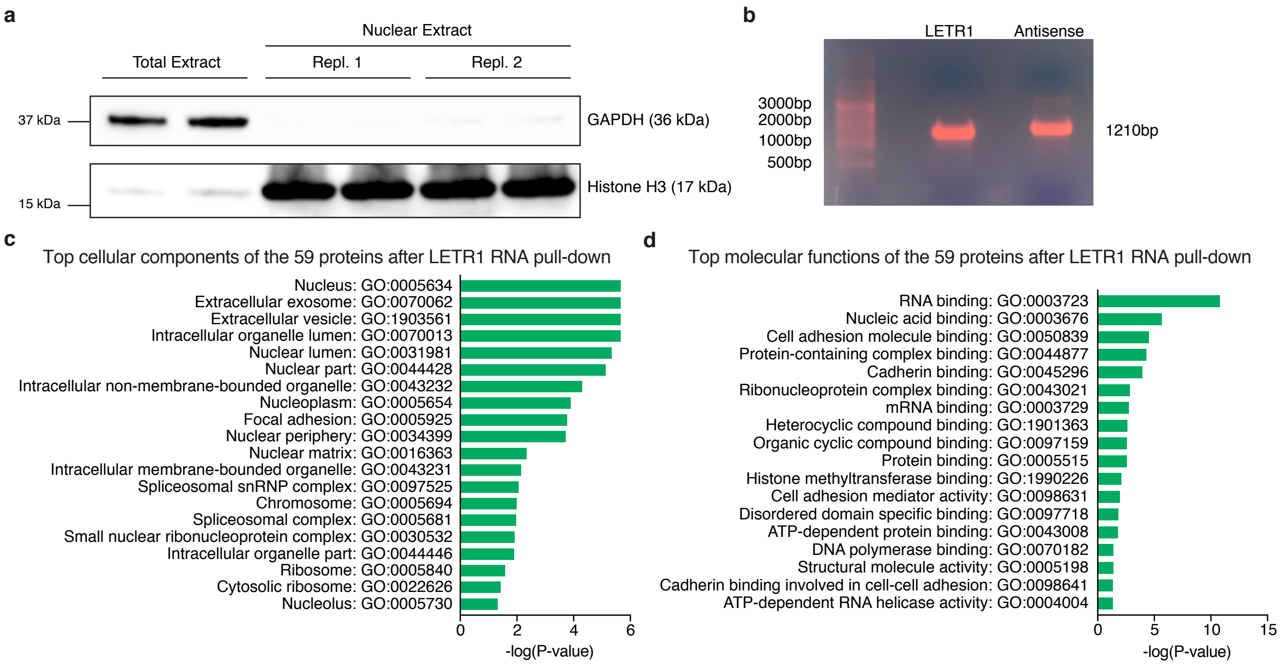

**Supplementary Figure 7: LETR1 RNA pull-down results.**

**(a)** Evaluation of the subcellular fractionation of nuclear compared to total extracts from neonatal LECs after performing western blot of GAPDH (cytoplasmic protein) and Histone H3 (nuclear protein). Uncropped western blot image is shown in Supplementary Figure 8.

**(b)** Gel electrophoresis showing the fragment size of biotin-LETR1 and antisense biotin-RNA control.

**(c, d)** Top significantly (P-value < 0.05) enriched GO terms for cellular components (c) and molecular functions (d) of the 59 proteins after LETR1 RNA pull-down, using g:ProfileR[2] (relative depth 4-8 for (c) and 1-4 for (d)).

Supplementary Figure 8

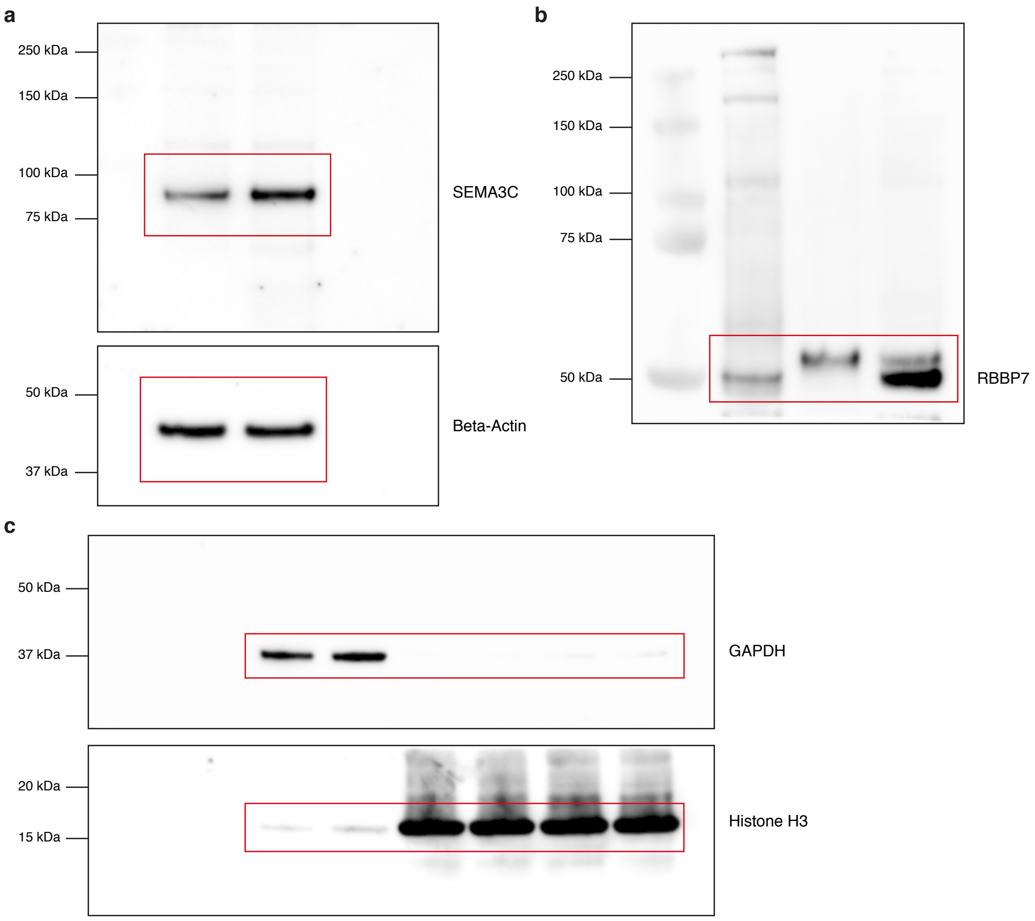

**Supplementary Figure 8: Uncropped western blot images.**

**(a)** Western blot images for SEMA3C from Supplementary Figure 6b.

**(b)** Western blot image for RBBP7 from Figure 7c.

**(c)** Western blot image for GAPDH and Histone H3 from Supplementary Figure 7a.

**References Supplemental Data Legends**
